# Identification of the main glutamine and glutamate transporters in *Staphylococcus aureus* and their impact on c-di-AMP production

**DOI:** 10.1101/754309

**Authors:** Merve S. Zeden, Igor Kviatkovski, Christopher F. Schuster, Vinai C. Thomas, Paul D. Fey, Angelika Gründling

## Abstract

A *Staphylococcus aureus* strain deleted for the c-di-AMP cyclase gene *dacA* is unable to survive in rich medium unless it acquires compensatory mutations. Previously identified mutations were in *opuD*, encoding the main glycine-betaine transporter, and *alsT*, encoding a predicted amino acid transporter. Here, we show that inactivation of OpuD restores the cell size of a *dacA* mutant to near wild-type size, while inactivation of AlsT does not, suggesting two different mechanisms for the growth rescue. AlsT was identified as an efficient glutamine transporter, indicating that preventing glutamine uptake in rich medium rescues the growth of the *S. aureus dacA* mutant. In addition, GltS was identified as a glutamine transporter. By performing growth curves with WT, *alsT* and *gltS* mutant strains in defined medium supplemented with ammonium, glutamine or glutamate, we revealed that ammonium and glutamine, but not glutamate promote the growth of *S. aureus*. This suggests that besides ammonium also glutamine can serve as a nitrogen source under these conditions. Ammonium and uptake of glutamine via AlsT inhibited c-di-AMP production, while glutamate uptake had no effect. These findings provide, besides the previously reported link between potassium and osmolyte uptake, a connection between nitrogen metabolism and c-di-AMP signalling in *S. aureus*.

**Graphical abstract:** 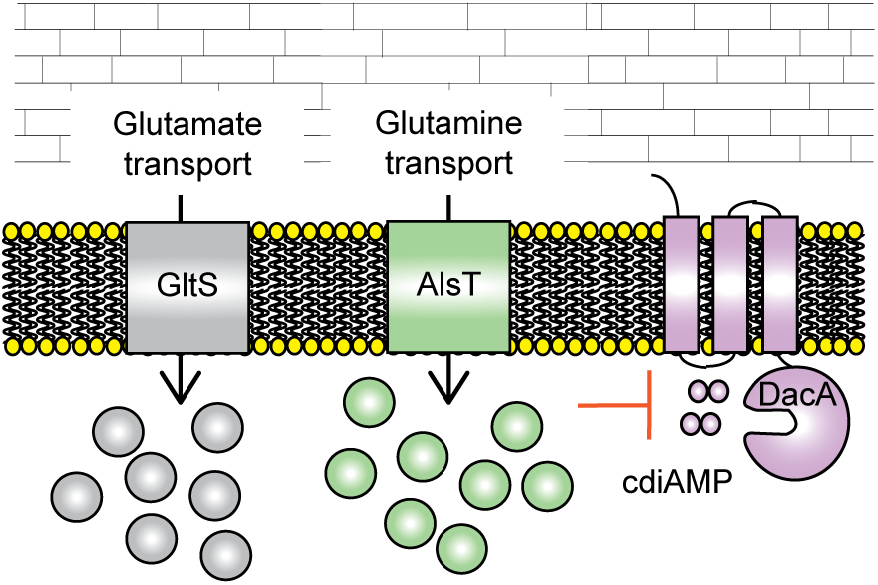

## Introduction

In the human host, S*taphylococcus aureus* can grow in virous tissues such as kidneys, bones, heart, soft tissues and lungs (Kluytmans *et al.*, 1997, Fridkin *et al.*, 2005). Sensitive regulatory mechanisms enable this organism to rapidly respond to external stimuli and environmental changes. Amongst others, this allows bacteria to adapt their metabolism and utilize different carbon and nitrogen sources available in each specific niche (Fridkin *et al.*, 2005, Spahich *et al.*, 2016, Vitko *et al.*, 2015, Crooke *et al.*, 2013, Fuller *et al.*, 2011, Richardson *et al.*, 2008, Halsey *et al.*, 2017, Lehman *et al.*, 2019).

Glucose is the preferred carbon source for *S. aureus,* but it can be limiting during infection due to the host immune response (Kelly & O’Neill, 2015, Spahich *et al.*, 2016, Halsey *et al.*, 2017, Lehman *et al.*, 2019). In glucose-limiting conditions, *S. aureus* instead catabolizes secondary carbon sources and amino acids, particularly glutamate and proline, which serve as major carbon sources during growth in the absence of glucose (Halsey *et al.*, 2017). However, not much is known about amino acid uptake and catabolism in *S. aureus* and how the availability of certain nutrients can affect virulence factor expression and invasion of the host. While a large number of amino acid transporters and oligopeptide permeases can be identified bioinformatically, their actual substrate specificities and functions in *S. aureus* have not yet been studied in detail. Predicting the substrates for transporters bioinformatically remains challenging and hence such questions need to be addressed experimentally.

Secondary messenger molecules are crucial in allowing bacteria to rapidly adapt to different environmental and host cell niches (Römling, 2008, Hengge, 2009). There is now considerable evidence that one of these messengers, cyclic di-adenosine monophosphate (c-di-AMP) plays a key role in osmoregulation in bacteria (Pham *et al.*, 2018, Pham & Turner, 2019, Quintana *et al.*, 2019, Zarrella *et al.*, 2018, Teh *et al.*, 2019, Fahmi *et al.*, 2019, Devaux *et al.*, 2018, Bai *et al.*, 2014, Zeden *et al.*, 2018, Corrigan *et al.*, 2011, Rocha *et al.*, 2019, Gundlach *et al.*, 2017a, Gundlach *et al.*, 2018, Witte *et al.*, 2013, Whiteley *et al.*, 2015, Whiteley *et al.*, 2017). c-di-AMP binds to and negatively regulates a number of different potassium and osmolyte importers (Rocha *et al.*, 2019, Quintana *et al.*, 2019, Kim *et al.*, 2015, Corrigan *et al.*, 2013, Moscoso *et al.*, 2015, Chin *et al.*, 2015, Huynh *et al.*, 2016, Schuster *et al.*, 2016, Pham & Turner, 2019, Pham *et al.*, 2018, Devaux *et al.*, 2018, Zarrella *et al.*, 2018, Gundlach *et al.*, 2017a, Gundlach *et al.*, 2018, Gundlach *et al.*, 2017b). In many Firmicutes, c-di-AMP is essential for bacterial growth under standard rich medium growth conditions, but it is also toxic at high levels. Hence, the cellular levels must be tightly regulated (Gundlach *et al.*, 2015, Mehne *et al.*, 2013, Corrigan *et al.*, 2011, Corrigan *et al.*, 2015, Woodward *et al.*, 2010, Witte *et al.*, 2013). In *S. aureus, Streptococcus agalactiae* and *Listeria monocytogenes,* deletion of *dacA* (also referred to as *cdaA* in many bacteria), coding for the diadenylate cyclase enzyme and responsible for the synthesis of c-di-AMP, was only possible in chemically defined medium (Whiteley *et al.*, 2015, Zeden *et al.*, 2018, Devaux *et al.*, 2018), whereas in *Bacillus subtilis* all three c-di-AMP cyclases could only be inactivated in minimal medium also containing low amounts of potassium (Gundlach *et al.*, 2017a).

Previously, we found that inactivation of the main glycine betaine transporter OpuD (SAUSA300_1245) as well as the predicted amino acid transporter AlsT (SAUSA300_1252) allows an *S. aureus dacA* mutant to grow in rich medium in the absence of c-di-AMP (Zeden *et al.*, 2018). In several other Firmicutes, including *B. subtilis, Lactococcus lactis, Streptococcus pneumoniae, S. agalactiae, Streptococcus pyogenes* and *L. monocytogenes*, inactivating mutations have also been identified in osmolyte and potassium transport systems that allow these bacteria to grow in the absence of c-di-AMP (Pham *et al.*, 2018, Pham & Turner, 2019, Quintana *et al.*, 2019, Zarrella *et al.*, 2018, Teh *et al.*, 2019, Fahmi *et al.*, 2019, Devaux *et al.*, 2018, Bai *et al.*, 2014, Zeden *et al.*, 2018, Corrigan *et al.*, 2011, Rocha *et al.*, 2019, Gundlach *et al.*, 2017a, Gundlach *et al.*, 2018, Witte *et al.*, 2013, Whiteley *et al.*, 2015, Whiteley *et al.*, 2017). This suggests that potassium and osmolyte transporters are more active in the absence of c-di-AMP, resulting in the accumulation of toxic levels of potassium and osmolytes in the cell. Consistent with a key function of c-di-AMP in regulating the osmotic balance in the cell, we found that *S. aureus* cells show significant differences in cell size depending on their intracellular c-di-AMP levels (Zeden *et al.*, 2018, Corrigan *et al.*, 2011). Cells of strain LAC\**gdpP,* which have high c-di-AMP levels, show a decrease in cell size, while cells of strain LAC\**dacA_G206S_*, containing low levels of c-di-AMP, show an increase in cell size (Zeden *et al.*, 2018, Corrigan *et al.*, 2011). As c-di-AMP negatively regulates potassium and osmolyte uptake (Corrigan *et al.*, 2013, Moscoso *et al.*, 2015, Schuster *et al.*, 2016), the increase in cell size is consistent with the hypothesis that an increase in potassium and osmolyte uptake and retention of water at reduced c-di-AMP levels leads to the observed increase in cell size. c-di-AMP levels affecting bacterial cell size has also been observed for other bacteria such as *S. pneumonia* and *L. monocytogenes* (Commichau *et al.*, 2019, Bai *et al.*, 2013).

AlsT is a predicted amino acid transporter and a correlation between cellular levels of c-di-AMP and the amino acids glutamate and glutamine has been reported for *B. subtilis* and *L. monocytogenes* (Whiteley *et al.*, 2017, Gundlach *et al.*, 2015, Gundlach *et al.*, 2018, Sureka *et al.*, 2014). A two-fold increase in cellular c-di-AMP levels was observed in *B. subtilis* when bacteria where grown in Spizizen minimal medium with glutamate (Glu) as compared to growth in the same medium with glutamine (Gln) as a nitrogen source (Gundlach *et al.*, 2015). In *L. monocytogenes*, c-di-AMP was identified as a negative regulator of the key TCA cycle enzyme pyruvate carboxylase (Sureka *et al.*, 2014). Depletion of c-di-AMP resulted in an increased flux into the TCA cycle and as a consequence an increase in the cellular glutamine/glutamate pool (Sureka *et al.*, 2014). In a *citZ* mutant, which lacks the TCA cycle enzyme citrate synthase and thus has an early block in the TCA cycle, the depletion of c-di-AMP no longer resulted in the accumulation of glutamate/glutamine in the cell (Sureka *et al.*, 2014). Interestingly, an *L. monocytogenes dacA*/*citZ* double mutant was again viable in rich medium (Sureka *et al.*, 2014, Whiteley *et al.*, 2017).

As part of this study, we further investigated why inactivation of the main glycine-betaine transporter OpuD and the predicted amino acid transporter AlsT allows *S. aureus* to grow in the absence of c-di-AMP in rich medium. We show that AlsT is a main glutamine transporter in *S. aureus* and that AlsT-mediated glutamine uptake represses c-di-AMP production. Similarly, growth in ammonium-containing defined medium but not in glutamate-containing medium, repressed c-di-AMP production. The repression of c-di-AMP production was independent of the activity of the c-di-AMP phosphodiesterase GdpP and the predicted cyclase regulator YbbR. With this study, we not only provide a further link between the c-di-AMP signalling network and osmotic regulation in bacterial cells but also with the uptake of specific nitrogen sources and amino acids in *S. aureus.*

## Results

### Inactivation of OpuD but not AlsT reduces the cell size of an *S. aureus dacA* mutant

In previous work, we reported a correlation between the cell size and c-di-AMP levels in *S. aureus*: bacteria with high c-di-AMP level are smaller, whereas bacteria with low c-di-AMP levels (strain LAC\**dacA_G206S_*) are larger as compared to wild-type bacteria (Zeden *et al.*, 2018, Corrigan *et al.*, 2011). We also reported that inactivating mutations in *opuD* (*SAUSA300_1245*) coding for the main glycine betaine osmolyte transporter and *alsT* (*SAUSA300_1252*) coding for a predicted amino acid sodium symporter, rescue the growth defect observed for the c-di-AMP negative *S. aureus* strain LAC\**dacA∷kan* in rich medium TSB (Zeden *et al.*, 2018). Here, we investigated further the mechanism by which the growth defect of the *dacA* mutant strain is rescued in the LAC\**dacA/opuD* and LAC\**dacA/alsT* suppressor strains. Initially, we compared the cell size of bacteria from the suppressor strains LAC\**dacA/opuD* and LAC\**dacA/alsT* to that of WT LAC* and the low c-di-AMP level strain LAC\**dacA_G206S_* after growth in the rich medium TSB. As expected, the bacteria with low-levels of c-di-AMP showed an increase in cell size as compared to WT bacteria (Fig. 1A and B). While a similar increase in cell size was still observed for bacteria of strain LAC\**dacA/alsT*, the cell size of LAC\**dacA/opuD* bacteria, while still increased as compared to the wild type, was significantly smaller as compared to the low-level LAC\**dacA_G206S_* strain (Fig. 1A and B). Because regular TSB medium is not suitable for the growth of the c-di-AMP null strain LAC\**dacA∷kan,* bacterial cell sizes were also determined following growth in TSB medium supplemented with 0.4 M NaCl, which is permissive for the growth of the *dacA* mutant (Fig. 1C-F). Similar to what was observed for the low c-di-AMP level *dacA_G206S_* mutant strain, the size of bacteria from the c-di-AMP null strain LAC\**dacA∷kan* was significantly increased compared to WT bacteria. As observed before, the cell size was not rescued for bacteria of the LAC\**dacA/alsT* suppressor strain (Fig. 1C-F). On the other hand, the size of LAC\**dacA/opuD* bacteria was similar to that of WT bacteria (Fig. 1C-F). Taken together, the observed differences in cell size indicate that the underlying molecular mechanisms enabling the *opuD* and *alsT* mutant strains to survive in the absence of c-di-AMP in rich medium might be different.

**Fig. 1.**
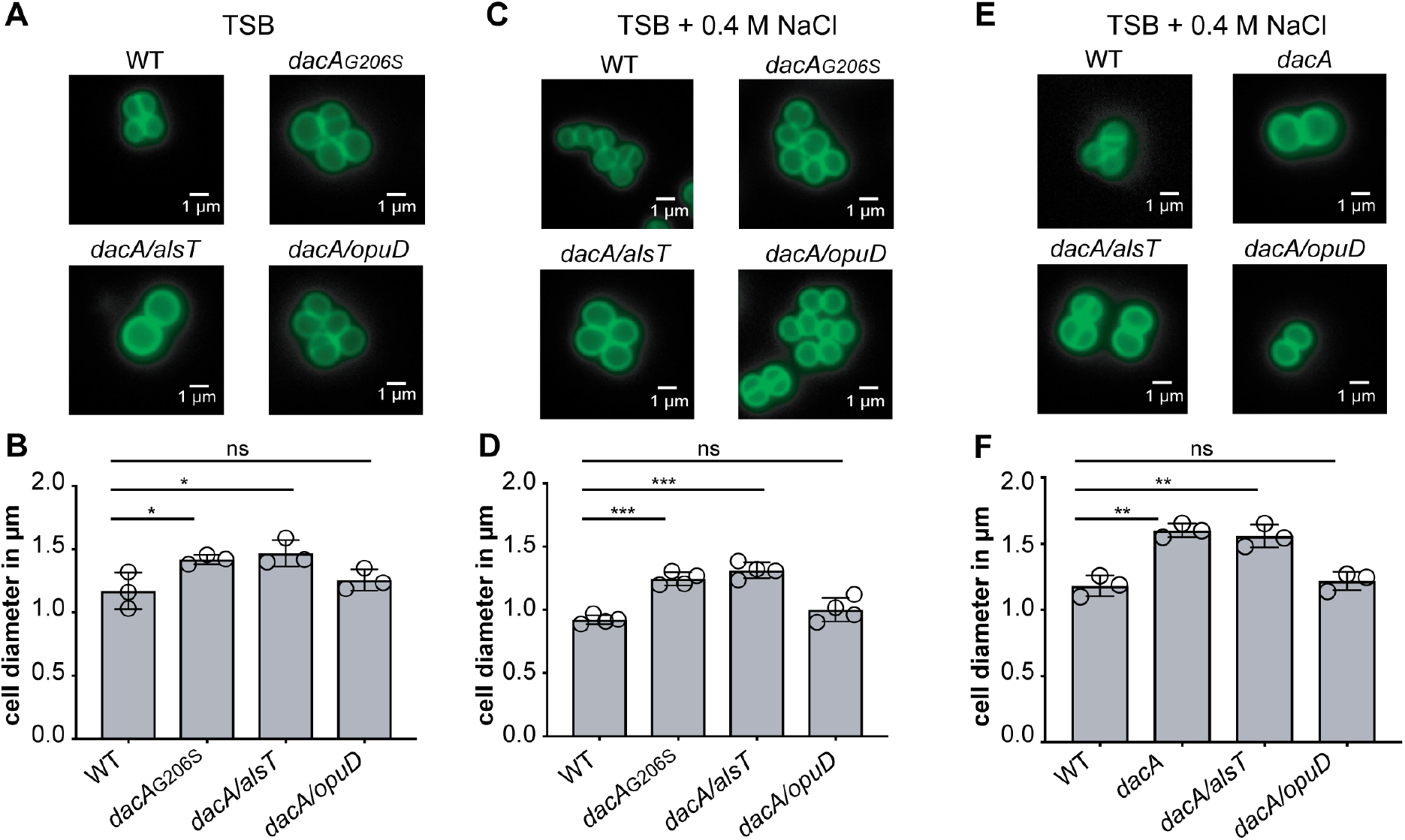
Inactivation of the glycine betaine transporter OpuD rescues the cell size of *S. aureus dacA* mutant bacteria. (A, C, E) Microscopy images of *S. aureus* cells stained with BODIPY-labelled vancomycin. Cultures of *S. aureus* LAC* (WT), LAC\**dacA*_G206S_ (*dacA*_G206S_) (panels A and C only), LAC*\*dacA∷kan* (*dacA*) (panel E only) and the suppressor strains LAC\**dacA/alsT* (*dacA/alsT*) and LAC\**dacA/opuD* (*dacA/opuD*) were grown in (A) TSB or (C and E) TSB 0.4 M NaCl medium and subsequently stained with fluorescently labelled vancomycin. The bacteria were then viewed using a fluorescent microscope and representative images are shown. Scale bars are 1 μm. (B, D, F) Bacterial cell diameter measurements. The diameter of non-dividing bacterial cells was measured as described in the Materials and Method section for *S. aureus* strains grown in (B) TSB or grown in (D and F) TSB 0.4 M NaCl medium. The diameters of 50 cells were determined and the average diameter calculated. The experiment was performed in triplicate (B and F) or quadruplicate (D) and the averages and SDs of the average cell diameters plotted. For statistical analysis, one-way ANOVAs followed by Dunnett’s multiple comparison tests were performed (ns = not significant, * = p<0.01, ** = p<0.001, *** = p<0.0001).

### AlsT is a glutamine transporter in *S. aureus*

AlsT (SAUSA300_1252) is a predicted amino acid transporter and annotated in the InterPro database (www.ebi.ac.uk/interpro) as alanine/sodium symporter. However, in a previous study no difference in the uptake of radiolabelled alanine was detected between a WT and the LAC\**dacA/alsT* mutant strain (Zeden *et al.*, 2018), indicating that AlsT is not an alanine transporter. To identify potential substrates for the *S. aureus* AlsT transporter, we initially followed the depletion of different amino acids from the culture supernatant during the growth of the WT LAC* strain and an isogenic *alsT* transposon mutant strain in TSB medium, where both strains exhibit similar growth rates (Fig. S1A). Of note, using this method, tryptophan uptake cannot be measured and it is also not possible to distinguish between glutamine/glutamate or asparagine/aspartate utilization. While no significant differences were observed for most amino acids (Fig. S1), a slight increase in the utilization of aspartate/asparagine and a slight decrease in the uptake of serine was observed (Fig. S1D and Fig. S1O), suggesting that AlsT could potentially be a serine transporter. To test this, uptake assays were performed with radiolabelled serine using WT LAC*, the *alsT* mutant strain LAC\**alsT∷tn* piTET as well as the complementation strain LAC\**alsT∷tn* piTET-*alsT*. However, no significant differences in the uptake rate of serine were observed between the strains (Fig. 2A), indicating that AlsT is not a main serine transporter in *S. aureus*. Next, a more detailed bioinformatics analysis was performed to identify potential AlsT substrates. A BlastP search against the *B. subtilis* 168 genome led to the identification of four proteins showing significant homology to the *S. aureus* AlsT (SAUSA300_1252) protein, namely AlsT (e-value: e-166), GlnT (e-value: e-149), YrbD (e-value: e-117) and YflA (e-value: 2e-72). Also in *S. aureus* a second AlsT homologue, SAUSA300_0914 (e-value 9e-108), could be identified (Fig. S2), which is encoded at a different chromosomal region. AlsT is annotated in *B. subtilis* as a potential glutamine sodium symporter (Zhu & Stülke, 2018), but to the best of our knowledge, this has not yet been experimentally verified. To test if *S. aureus* AlsT is a potential glutamine or glutamate transporter, uptake assays were performed with radiolabelled glutamine and glutamate using the WT *S. aureus* strain LAC*, the *alsT* mutant LAC\**alsT∷tn* piTET and the complementation strain LAC\**alsT∷tn* piTET-*alsT*. Uptake of glutamine, but not of glutamate, was severely reduced in the *alsT* mutant when compared to the WT strain (Fig. 2B-C). This defect was restored upon expression of *alsT* in the complementation strain (Fig. 2C). To confirm that *alsT* functions as main glutamine transporter also in the LAC\**dacA/alsT* suppressor strain, uptake assays were also performed with strain LAC\**dacA/alsT* along with the WT LAC* and LAC\**dacA∷kan* control strains (Fig. 2D-F). Similar as observed for the *alsT* single mutant, serine and glutamate uptake were only marginally affected in strain LAC\**dacA/alsT* (Fig. 2D-E), whereas glutamine uptake was severely reduced in strain LAC\**dacA/alsT* when compared to the control strains (Fig. 2F). These data suggest that under the uptake assay conditions tested, AlsT functions as the main glutamine transporter in *S. aureus.* A slight reduction in glutamine and serine uptake was seen in the absence of c-di-AMP (Fig. 2D and Fig. 2F), suggesting that c-di-AMP levels can impact glutamine and serine uptake in *S. aureus.* Taken together, our data suggest that *S. aureus* cells that are unable to produce c-di-AMP can survive in rich medium such as TSB, when glutamine uptake is reduced or blocked. However, we cannot formally exclude that AlsT is able to transport other amino acid or substrates present in rich medium.

**Fig. 2.**
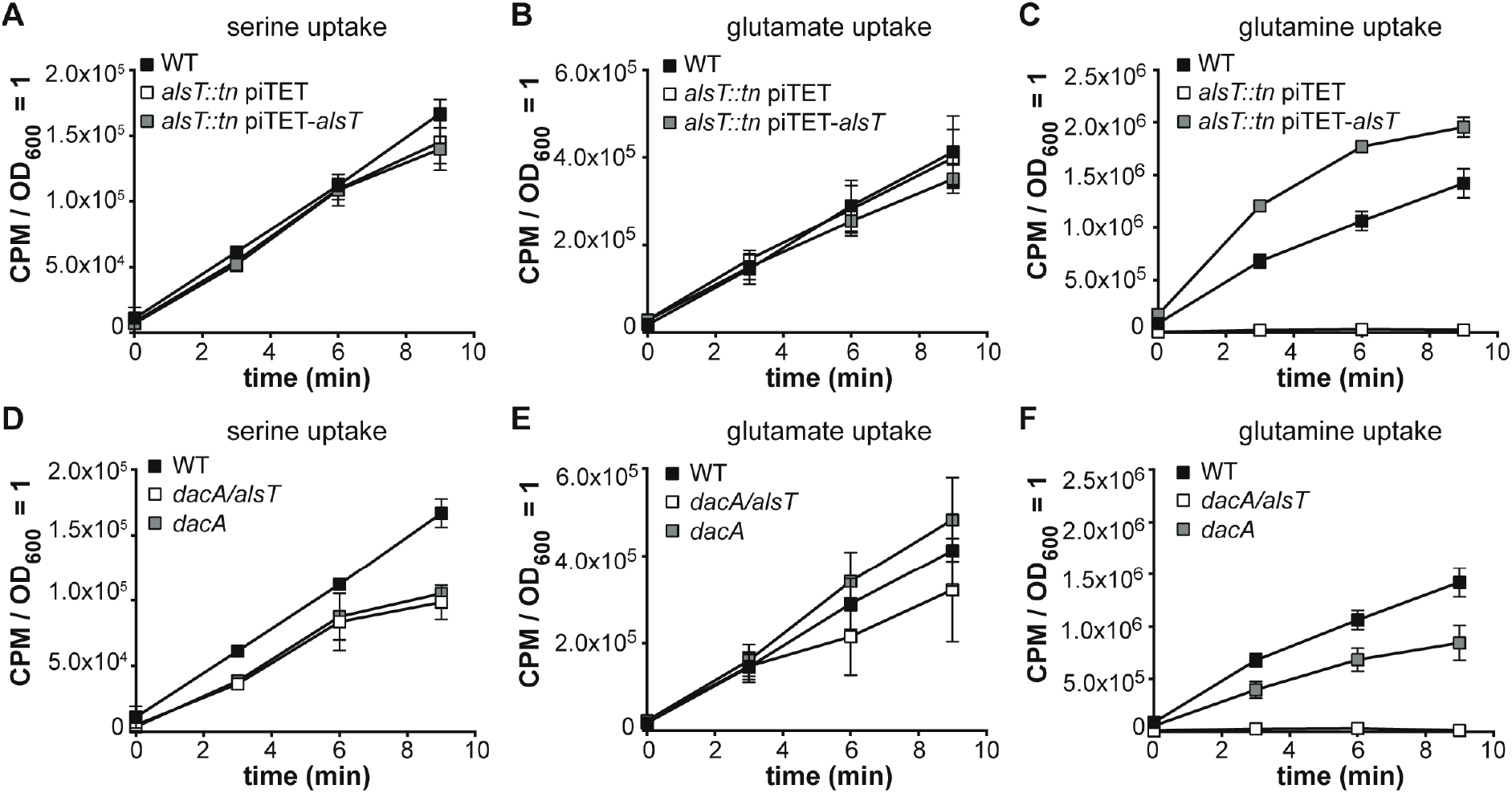
AlsT is a glutamine transporter in *S. aureus*. (A-F) Amino acid uptake assays. (A-C) *S. aureus* strain LAC* (WT), the *alsT* mutant LAC\**alsT∷tn* piTET (*alsT∷tn* piTET) and the complementation strain LAC\**alsT∷tn* piTET-*alsT* (*alsT∷tn* piTET-*alsT*) were grown to mid-log phase in glucose defined media as indicated in the method section and supplemented with 200 ng/ml Atet for the strains containing plasmids. Subsequently radiolabelled (A) serine, (B) glutamate or (C) glutamine was added to culture aliquots, samples removed and filtered at the indicated time points and the radioactivity accumulated in the cells measured. The average values and SDs from three (A, C, D, F) or four (B, E) experiments were plotted. (D-F) The same uptake assay experiment was performed as described in (A-C) but using *S. aureus* strains LAC\**dacA∷kan* (*dacA*) and LAC\**dacA/alsT* (*dacA/alsT*). The amino acid uptake curve for the LAC* (WT) strain is the same as shown in panels A-C, as all strains were grown and processed at the same time.

### Investigating the contribution of SAUSA300_0914 and GlnQ to glutamine and glutamate transport in *S. aureus*

*S. aureus SAUSA300_0914* codes for a predicted amino acid symporter, which shows 41% identity to the *S. aureus* AlsT protein. After assigning AlsT a function as glutamine transporter, we wanted to test if SAUSA300_0914 might also play a role in glutamine or glutamate transport. To this end, strain LAC\**0914∷tn* was constructed by transducing the genomic region from the NMTL strain NE1463 (Fey *et al.*, 2013) containing a transposon insertion in *SAUSA300_0914* into the *S. aureus* LAC* background. Subsequently the uptake of radiolabelled glutamine and glutamate was assessed (Fig. 3A-B). No significant differences in the uptake of these amino acids was observed between WT LAC* and strain LAC\**0914∷tn*, showing that SAUSA300_0914 does not function as a major glutamine or glutamate transporter under our assay conditions (Fig. 3A-B). AlsT and SAUSA300_0914 are members of the amino acid/sodium symporter family of transporters, which are composed of a single multimembrane spanning protein. Besides this type of transporter, GlnPQ-type ABC transporters play a major role in glutamine and glutamate transport in other bacteria (Schuurman-Wolters & Poolman, 2005). *S. aureus* contains a *glnPQ (SAUSA300_1808 - SAUSA300_1807)* operon with *glnP* coding for a substrate binding domain-permease fusion protein and *glnQ* coding for the cytoplasmic nucleotide-binding ATPase domain. The results from a previous study suggested that this transporter functions as glutamine transporter in *S. aureus*, as a *glnP* mutant was more resistant to the toxic glutamine analogue γ-L-glutamyl hydrazide (Zhu *et al.*, 2009). To assess the contribution of the GlnPQ transporter to glutamine and glutamate transport in *S. aureus* LAC*, the strain LAC\**glnQ∷tn* was generated by transducing the *glnQ∷tn* region from the NMTL strain NE153 (Fey *et al.*, 2013) into the LAC* background. The resulting *glnQ* mutant strain LAC\**glnQ∷tn* displayed no difference in glutamine or glutamate uptake compared to WT LAC* (Fig. 3C-D). This indicates that the ABC transporter GlnPQ does not function under our assay conditions and in the *S. aureus* LAC* strain background as a main glutamate or glutamine transporter.

**Fig. 3.**
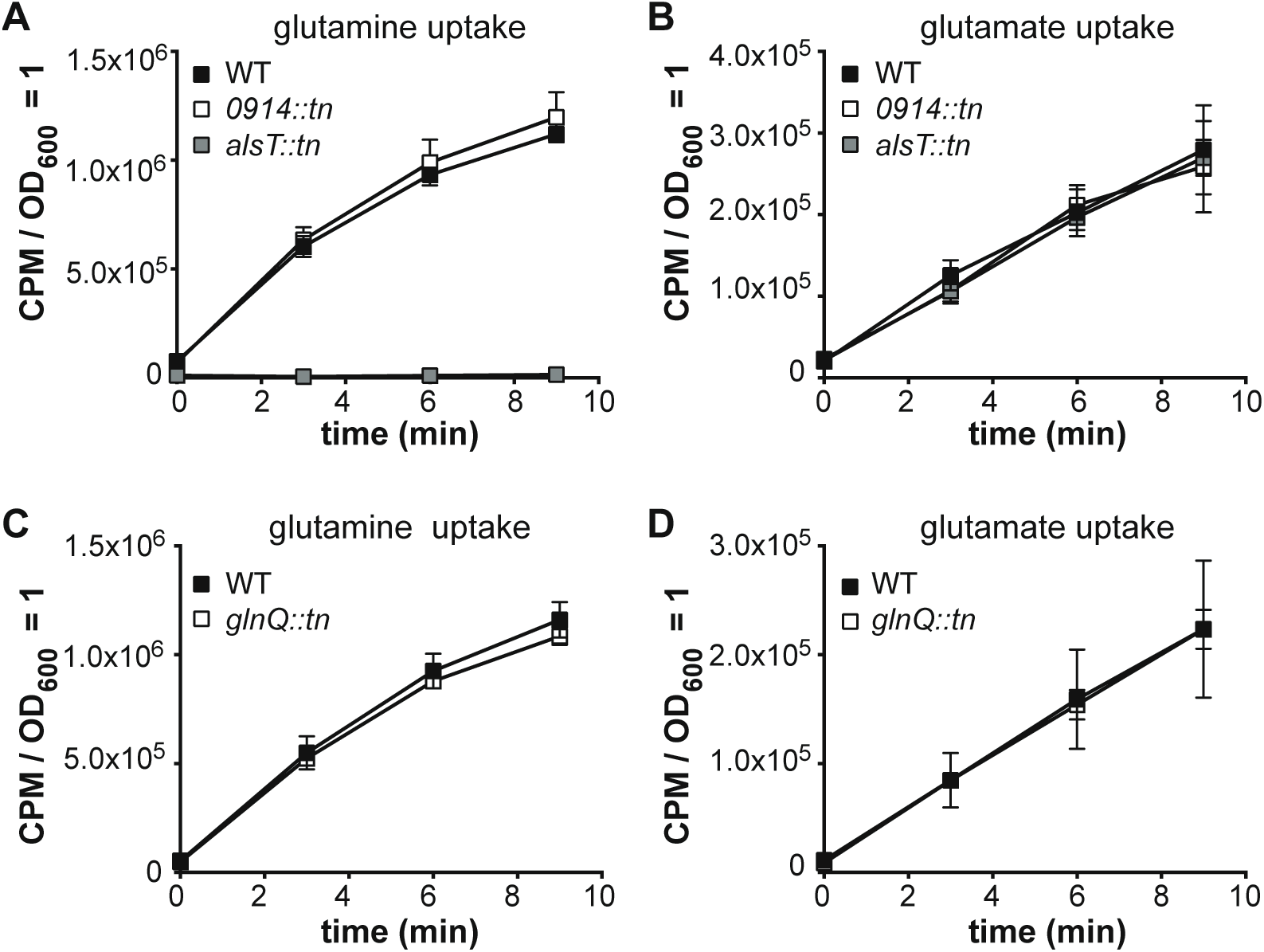
LAC* *0914∷tn* and LAC* *glnQ∷tn* strains do not show a defect in glutamine or glutamate uptake. Amino acid uptake assays. (A and B) *S. aureus* strains LAC* (WT), LAC\**0914∷tn* and LAC\**alsT∷tn* were grown to mid-log phase in GDM+Glu+NH_3_. Subsequently radiolabelled (A) glutamine or (B) glutamate was added to culture aliquots, samples removed and filtered at the indicated time points and the radioactivity accumulated in the cells measured. The average values and SDs from three experiments were plotted. (C and D) Amino acid uptake assays were performed and the data plotted as described in panels A and B, but using *S. aureus* strains LAC* (WT) and LAC\**glnQ∷tn* (*glnQ∷tn*).

### Inactivation of AlsT but not SAUSA300_0914 or GlnQ reduces the susceptibility of *S. aureus* to the toxic glutamine analogue γ-L-glutamyl hydrazide

To further validate the findings from the uptake assays and verify that AlsT is the main glutamine transporter, we performed growth curves in the presence of increasing concentrations of the toxic glutamine analogue γ-L-glutamyl hydrazide with the WT and LAC\**alsT∷tn* mutant strains. Strains LAC\**0914∷tn* and LAC\**glnQ∷tn* were also included in these assays, to uncover a potential low level glutamine uptake activity for the SAUSA300_0914 and GlnPQ transporters. Strains defective in taking up this glutamine analogue are expected to show reduced susceptibility to this toxic compound. In the absence of the compound, all strains grew similarly in the chemically defined medium used for this assay (Fig. 4A). As expected, addition of γ-L-glutamyl hydrazide reduced the growth of the WT LAC* strain, in a dose-deponent manner (Fig. 4B and Fig. S3). Similar growth inhibition curves to that of the WT strain were obtained for strains LAC\**0914∷tn* and LAC\**glnQ∷tn*, while strain LAC\**alsT∷tn* showed increased resistance to the compound (Fig. 4B and Fig. S3). These findings support our earlier conclusion that AlsT is the main glutamine transporter in *S. aureus*, while GlnPQ and SAUSA300_0914 are either unable to take up glutamine or play only a minor role in its uptake under our growth conditions.

**Fig. 4.**
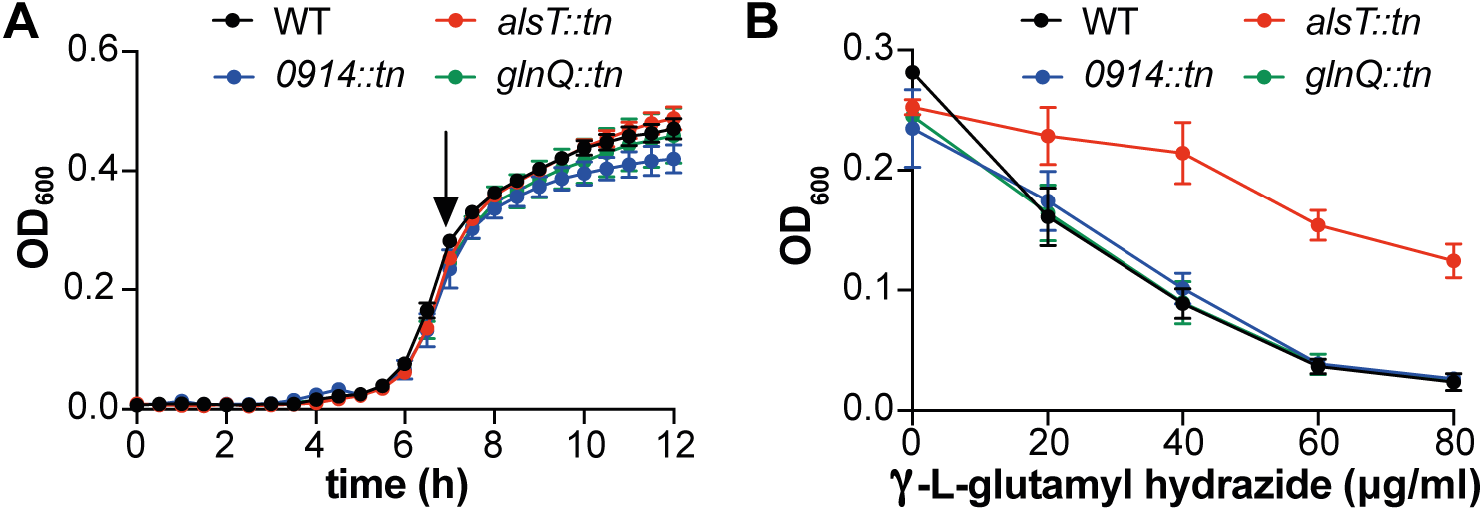
LAC* *alsT∷tn* shows increased resistance to the toxic glutamine analogue γ-L-glutamyl hydrazide. (A) Bacterial growth curves. *S. aureus* strains LAC* (WT), LAC\**alsT∷tn*, LAC\**0914∷tn* and LAC\**glnQ∷tn* were grown in GDM+NH_3_ medium in 96-well plates and OD_600_ reading determined over 12 h. The average OD_600_ values and SDs from three independent biological replicates were plotted. (B) γ-L-glutamyl hydrazide susceptibility assay. The same *S. aureus* strains as in (A) were grown in GDM+NH_3_ medium in the absence or presence of γ-L-glutamyl hydrazide at a final concentration of 20, 40, 60 or 80 μg/ml. OD_600_ reading were determined over 12 h and the complete growth curves are shown in Figure S3. In this graph, the average OD_600_ values from the 7 h time point (marked with an arrow in (A)) and SDs from three biological replicates were plotted against the γ-L-glutamyl hydrazide concentration in the growth medium.

### GltS (SAUSA300_2291) is a glutamate transporter in *S. aureus*

*S. aureus* does not only take up glutamine but also shows robust glutamate uptake (Fig. 2 and Fig. 3). However, none of the transporters (AlsT, SAUSA300_0914 and GlnPQ) investigated so far plays a major role in glutamate uptake under our growth conditions. In *B. subtilis* GltT, belonging to the dicarboxylate/amino acid cation symporter (DAACS) family of proteins, is a major high-affinity Na^+^-coupled glutamate/aspartate symporter and can also mediate the uptake of glyphosate (Wicke *et al.*, 2019). Two paralogs, DctP and GltP, are found in *B. subtilis* of which GltP has also been shown to be a glutamate transporter (Tolner *et al.*, 1995). The *S. aureus* protein SAUSA300_2329 (from here on referred to as GltT) shows a high degree of similarity (52% identity) to the *B. subtilis* GltT protein. In addition, SAUSA300_2291 (from here on referred to as GltS) is annotated in UniProt (www.uniprot.org) as a potential glutamate transporter in *S. aureus*. To experimentally test if GltT or GltS impact glutamate transport in *S. aureus*, strains LAC\**gltT∷tn* and LAC\**gltS∷tn* were constructed by moving the respective *gltT and gltS* transposon insertion regions from the NMTL strains NE566 and NE560 (Fey *et al.*, 2013) into the LAC* strain background. Next, the uptake of radiolabelled glutamine and glutamate was assessed for WT LAC* and strains LAC* *gltT∷tn* and LAC* *gltS∷tn*. No difference in the uptake of glutamine was observed between the strains (Fig. 5A) and in the case of LAC\**gltT∷tn*, also no difference in the uptake of glutamate was observed (Fig. 5B). However, a significant reduction in glutamate uptake was observed for strain LAC\**gltS∷tn* when compared to the WT strain (Fig. 5B). The glutamate uptake defect could be restored in a complementation strain harbouring plasmid piTET-*gltS* allowing for inducible *gltS* expression (Fig. 5C). Indeed, increased glutamate uptake was observed in the complementation strain, indicating increased *gltS* expression in the complementation strain as compared to the WT strain. Taken together, these data indicate that under the growth conditions tested, GltS is the main glutamate transporter in *S. aureus*.

**Fig. 5.**
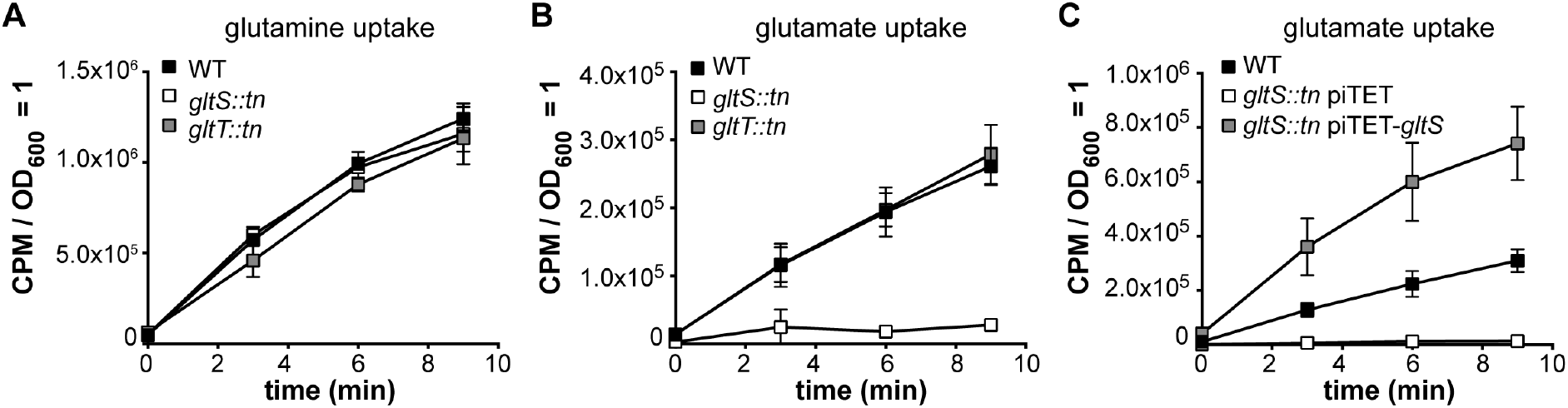
GltS is a glutamate transporter in *S. aureus*. Amino acid uptake assays. (A and B) *S. aureus* strains LAC* (WT), LAC\**gltT∷tn* and LAC\**gltS∷tn* were grown to mid-log phase in GDM+Glu+NH_3_. Subsequently radiolabelled (A) glutamine or (B) glutamate was added to culture aliquots, samples removed and filtered at the indicated time points and the radioactivity accumulated in the cells measured. (C) Same as (B) but using *S. aureus* strains LAC* (WT), LAC\**gltS∷tn* piTET and the complementation strain LAC\**gltS∷tn* piTET-*gltS* and supplementing the GDM+Glu+NH_3_ medium with 200 ng/μl Atet. The average values and SDs from three experiments were plotted.

### Ammonium and glutamine but not glutamate stimulate the growth of *S. aureus* in defined medium containing glucose as carbon source

Glutamine and glutamate are important amino acids that can serve, together with ammonium, as nitrogen sources for the synthesis of many other cellular metabolites. Since *S. aureus* is phenotypically auxotroph for many amino acids and at the same time can use several amino acids as carbon and nitrogen sources, it is not possible to grow this organism in any of the typical minimal media that are used to assess the ability of bacteria to specifically use ammonium, glutamine or glutamate as nitrogen sources. However, to begin to examine the effect of these compounds on the growth of *S. aureus*, growth curves were performed with the WT LAC* strain in glucose containing defined medium (GDM), containing essential vitamins, metals and 17 amino acids but lacking ammonium, glutamine and glutamate as potential nitrogen/amino acid sources (see Table S1 for medium composition). In addition, the WT LAC* was also grown in GDM containing glutamine (GDM+Gln), glutamate (GDM+Glu), ammonium (GDM+NH_3_), glutamine and ammonium (GDM+Gln+NH_3_) or glutamate and ammonium (GDM+Glu+NH_3_). The addition of glutamine or ammonium alone or in combinations stimulated the growth of the WT LAC* strain as compared to its growth in GDM (Fig. 6A). On the other hand, no growth improvement was seen in the presence of glutamate (GDM+Glu) (Fig. 6A). To examine the contribution of the glutamine and glutamate transporters AlsT and GltS, additional growth curves were performed in the different media with WT LAC* as well as strains LAC\**alsT∷tn* and LAC\**gltS∷tn* (Fig. S4). Similar growth profiles were observed for all strains in the different media (Fig. S4), except in GDM+Gln, in which the *alsT* mutant strain exhibited reduced growth compared to the WT and *gltS* mutant strains (Fig. S4B). The growth defect could be restored in the *alsT* complementation strain harbouring plasmid piTET-*alsT* (Fig. 6B). Taken together, these data indicate that ammonium and glutamine are preferred over glutamate for the growth of *S. aureus*. The observation that the addition of ammonium improves the growth of *S. aureus* indicates that our base medium is likely nitrogen limiting and suggests that glutamine but not glutamate can likely also serve as nitrogen source under these growth conditions. Finally, these data further confirm the importance of AlsT for glutamine uptake in *S. aureus.*

**Fig. 6.**
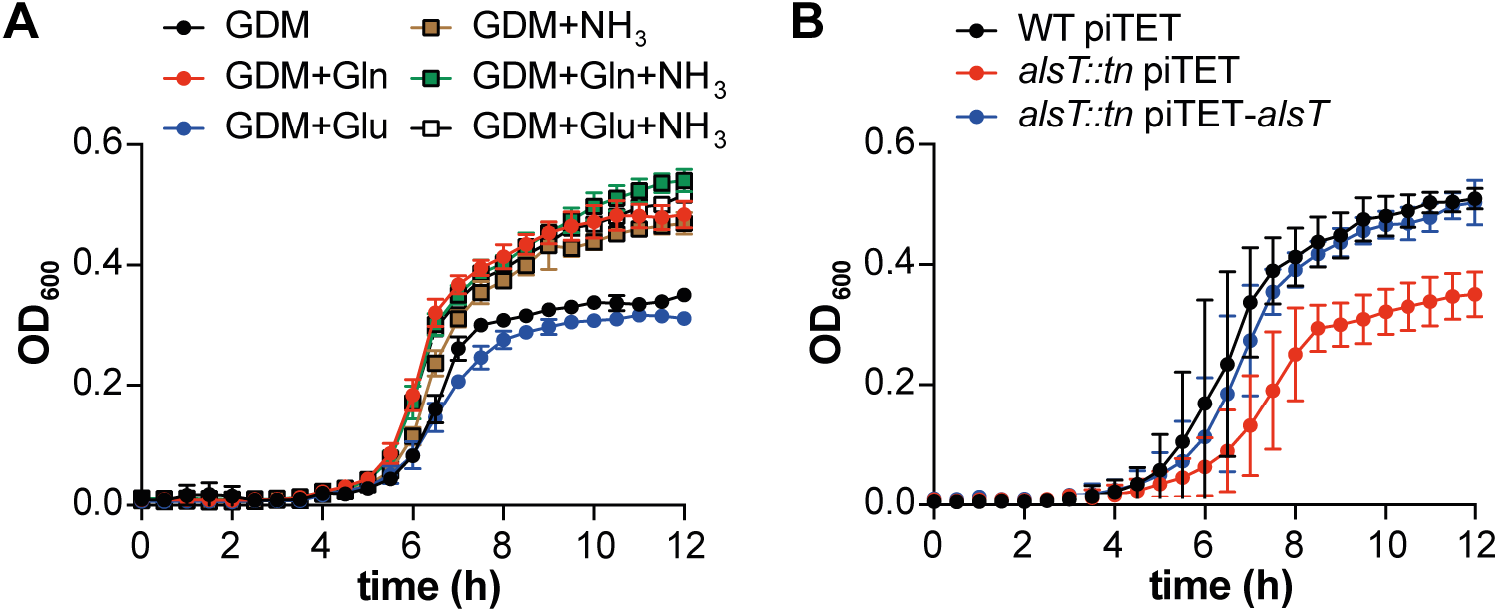
Addition of glutamine and ammonium but not glutamate stimulates the growth of *S. aureus* in glucose-containing defined medium. (A) Bacterial growth curves. The WT *S. aureus* strain LAC* was grown in 96-well plates in glucose-containing defined medium (GDM) or in GDM supplemented with glutamine (Gln), glutamate (Glu), ammonium (NH_3_) or combinations thereof as specified in the legend. OD_600_ readings were determined every 30 min and the average and SDs of three biological replicates plotted. (B) Bacterial growth curves. *S. aureus* strains LAC* piTET (WT piTET), the *alsT* mutant strain LAC\**alsT∷tn* piTET and the complementation strain LAC\**alsT∷tn* piTET-*alsT* were grown in GDM+Gln medium supplemented with 200 ng/μl Atet. OD_600_ readings were determined every 30 min and the average and SDS of three biological replicates plotted.

### Ammonium and glutamine uptake lead to a reduction in c-di-AMP levels in *S. aureus*

For *B. subtilis,* it has been reported that the addition of glutamine, glutamate or ammonium to a defined growth medium can affect cellular c-di-AMP levels (Gundlach *et al.*, 2015). It was further proposed that glutamate uptake and to some extent also ammonium uptake leads to an activation of c-di-AMP synthesis in this organism (Gundlach *et al.*, 2015). To assess if the presence of glutamine, glutamate or ammonium would also affect c-di-AMP levels in *S. aureus*, the intracellular c-di-AMP concentrations were determined for the WT *S. aureus* strain LAC* following growth in GDM, GDM+Gln, GDM+Glu, GDM+NH_3_, GDM+Gln+NH_3_ and GDM+Glu+NH_3_. Using a competitive ELISA assay, c-di-AMP could be readily detected in bacteria grown in GDM, our base medium (Fig. 7A). Similar amounts of c-di-AMP were detected in bacteria grown in the glutamate-containing medium (GDM+Glu), however the c-di-AMP levels were significantly lower in bacteria grown in medium containing either ammonium or glutamine (GDM+Gln, GDM+NH_3_, GDM+Gln+NH_3_, GDM+Glu+NH_3_) (Fig. 7A). To verify that the addition of glutamine reduces c-di-AMP production and to investigate the contribution of the glutamine transporter AlsT to this inhibition, c-di-AMP levels were determined for WT piTET, the *alsT* mutant LAC\**alsT∷tn* piTET and the complementation strain LAC\**alsT∷tn* piTET-*alsT* following growth in the glutamine containing medium GDM+Gln. While again low c-di-AMP levels were detected for the WT strain, the c-di-AMP levels increased significantly in the *alsT* mutant and were restored back to wild-type levels in the complementation strain (Fig. 7B). A similar experiment was performed with the *gltS* mutant and complementation strain in the glutamate containing medium GDM+Glu. High and similar c-di-AMP levels were detected for all strains (Fig. 7C), indicating that neither the addition of glutamate to the medium nor its uptake impacts c-di-AMP production in *S. aureus* under our test conditions. Taken together, these data highlight that ammonium as well as AlsT-mediated glutamine uptake represses c-di-AMP production in *S. aureus*.

**Fig. 7.**
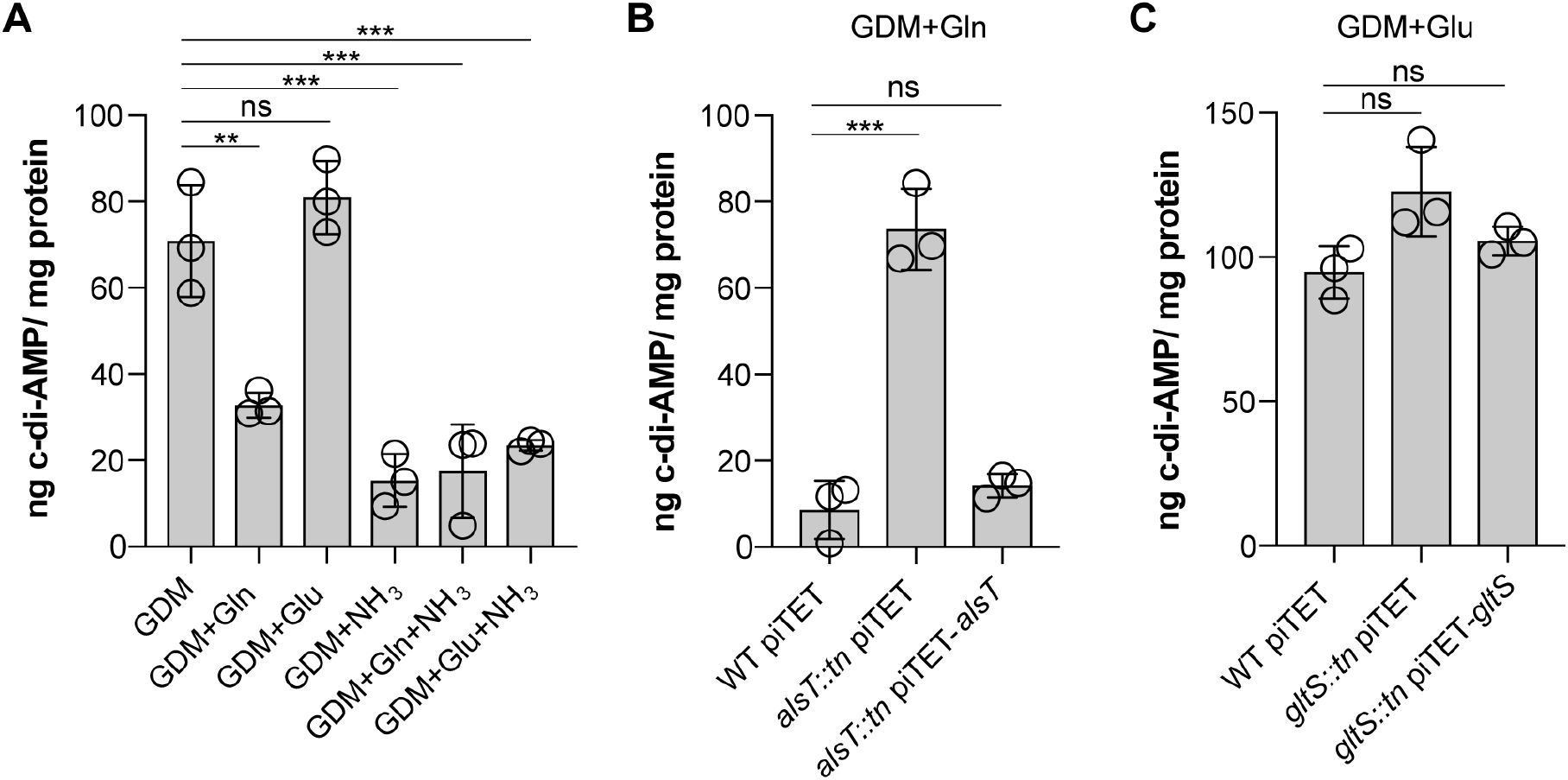
Ammonium and glutamine uptake inhibit c-di-AMP production in *S. aureus*. (A-C) Cellular c-di-AMP levels. (A) The WT *S. aureus* strain LAC* was grown in GDM or GDM supplemented with glutamine (Gln), glutamate (Glu), ammonium (NH_3_), or combinations thereof as indicated on the X-axes. Cell extracts were prepared and c-di-AMP concentrations measured using a competitive ELISA assay. The average values and SDs from three biological replicates were plotted as ng c-di-AMP/mg protein. For statistical analysis, one-way ANOVAs followed by Dunnett’s multiple comparison tests were performed to identify statistically significant differences between the different media as compared to GDM (ns = not significant, ** = p<0.001, *** = p<0.0001). (B) *S. aureus* strains LAC* piTET (WT piTET), LAC\**alsT∷tn* piTET and the complementation strain LAC\**alsT∷tn* piTET-*alsT* were grown in GDM+Gln supplemented with 200 ng/μl Atet and c-di-AMP levels determined and plotted as described in (A). (C) *S. aureus* strains LAC* piTET (WT piTET), LAC\**gltS∷tn* piTET and the complementation strain LAC\**gltS∷tn* piTET-*gltS* were grown in GDM+Glu supplemented with 200 ng/μl Atet and c-di-AMP levels determined and plotted as described in (A).

### The inhibition of the c-di-AMP production by glutamine and ammonium is not mediated by GdpP or YbbR

The observed reduction of c-di-AMP levels in the presence of glutamine or ammonium could potentially be achieved through an increase in the activity of the c-di-AMP specific phosphodiesterase GdpP. To investigate this, cellular c-di-AMP levels were compared between the WT LAC* and the isogenic *gdpP* mutant strain LAC\**gdpP∷kan*. As previously reported for strain LAC\**gdpP∷kan* following growth in TSB medium (Corrigan *et al.*, 2011), the c-di-AMP levels were also increased in the *gdpP* mutant compared to the WT strain following growth in GDM, the glucose containing defined medium used as part of this study (Fig. 8A). However, a significant reduction in the cellular c-di-AMP levels was also seen for the *gdpP* mutant following the addition of glutamine or ammonium to the medium (Fig. 8A). This indicates that the reduction in c-di-AMP levels upon addition of glutamine or ammonium is likely due to decreased synthesis by DacA and not increased degradation by GdpP. We next tested the involvement of YbbR, a proposed c-di-AMP cyclase regulator, by comparing the cellular c-di-AMP levels produced by WT LAC* and strain LAC*Δ*ybbR*. Similar c-di-AMP levels were detected in the WT and *ybbR* mutant in GDM medium (Fig. 8B). The addition of glutamine or ammonium to the medium led also to a large reduction in the cellular c-di-AMP in the *ybbR* mutant strain (Fig. 8B). These data suggest that the observed reduction of c-di-AMP production in the presence of glutamine and ammonium is neither mediated by GdpP nor YbbR, and hence involves a different regulator protein, or that the cellular glutamine and nitrogen levels are directly sensed by the cyclase DacA.

**Fig. 8.**
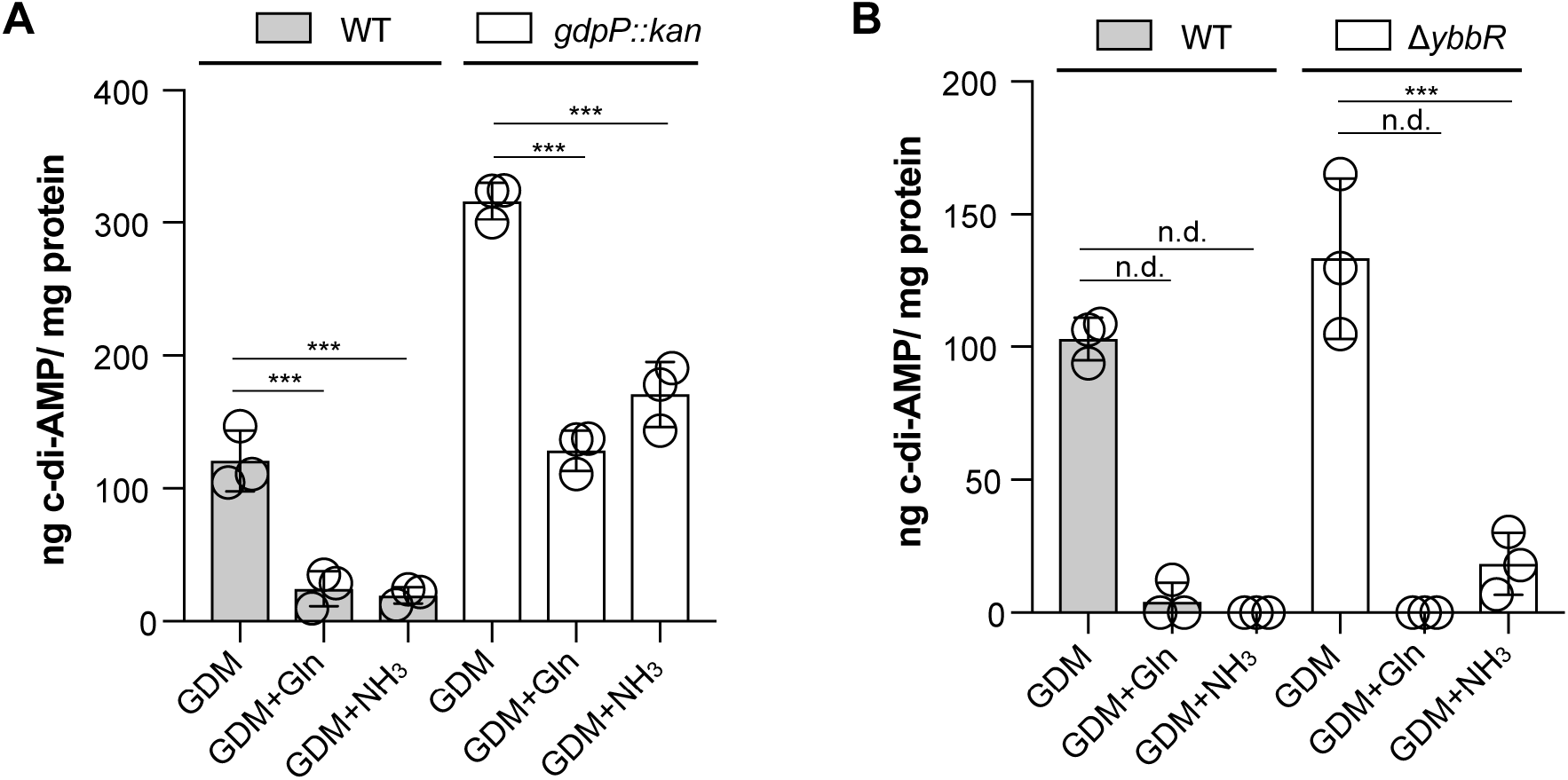
The inhibition of c-di-AMP production by glutamine and ammonium is independent of GdpP and YbbR. (A and B) Cellular c-di-AMP levels. (A) *S. aureus* strains LAC* (WT) and LAC\**gdpP∷kan* (*gdpP∷kan)* were grown in GDM or in GDM containing glutamine (GDM+Gln) or ammonium (GDM+NH_3_). Cell extracts were prepared, and c-di-AMP concentrations measured using a competitive ELISA assay. The average values and SDs from three biological replicates were plotted as ng c-di-AMP/ mg protein. For statistical analysis one-way ANOVAs followed by Dunnett’s multiple comparison tests were performed to identify statistically significant differences between the different media as compared to the GDM medium (n.d. = not determined. Statistical analysis was not performed due to values below our detection limit; these values were set to 0, ** = p<0.001, *** = p<0.0001). (B) Same as in (A) but using *S. aureus* strains LAC* (WT) and LAC*Δ*ybbR* (Δ*ybbR*).

## Discussion

Over the last decade, considerable evidence has emerged that c-di-AMP plays a major role in osmotic regulation in bacteria, primarily by positively regulating potassium export or negatively regulating potassium and osmolyte uptake (Rocha *et al.*, 2019, Quintana *et al.*, 2019, Kim *et al.*, 2015, Corrigan *et al.*, 2013, Moscoso *et al.*, 2015, Chin *et al.*, 2015, Huynh *et al.*, 2016, Schuster *et al.*, 2016, Pham & Turner, 2019, Pham *et al.*, 2018, Devaux *et al.*, 2018, Zarrella *et al.*, 2018, Gundlach *et al.*, 2017a, Gundlach *et al.*, 2018, Gundlach *et al.*, 2017b, Gundlach *et al.*, 2019). However, individual c-di-AMP target proteins identified thus far are themselves not essential. Therefore, the essentiality of c-di-AMP is likely due to its ability to regulate multiple target proteins simultaneously. Furthermore, in the absence of this molecule, many transporters are activated rather than inactivated, likely leading to accumulation of toxic levels of metabolites, such as potassium and osmolytes. Consistent with this idea, inactivating mutations in potassium uptake systems, oligopeptide and osmolyte transporters have been reported to rescue the growth defect of bacteria unable to produce c-di-AMP (Whiteley *et al.*, 2015, Whiteley *et al.*, 2017, Gundlach *et al.*, 2017a, Gundlach *et al.*, 2017b, Pham *et al.*, 2018, Devaux *et al.*, 2018, Zeden *et al.*, 2018). We have previously shown that inactivation of the main glycine betaine transporter OpuD bypasses the requirement of c-di-AMP for the growth of *S. aureus* in rich medium (Zeden *et al.*, 2018). We hypothesize that inactivation of OpuD might help a c-di-AMP null strain survive by allowing bacteria to re-establish their osmotic balance. Bacteria of the *dacA/opuD* mutant strain, which cannot produce c-di-AMP but are also defective in glycine betaine transport, are similar in size to WT bacteria (Fig. 1). At this point it is not known if c-di-AMP can directly bind to and regulate the activity of the *S. aureus* OpuD protein. We attempted to address this question; however, despite using multiple different approaches, we were unable to produce sufficient amounts of the full-length OpuD membrane protein to perform protein/nucleotide interaction studies. On the other hand, a direct role for c-di-AMP in the control of glycine betaine or betaine transporters has been reported in *S. agalactiae* and *L. lactis*. In these organisms, c-di-AMP binds to the transcriptional regulator BusR, which controls the expression of the predicted glycine betaine or betaine transporter BusAB (Devaux *et al.*, 2018, Pham *et al.*, 2018).

Bacteria of the *dacA/alsT* suppressor strain, which survive in the absence of c-di-AMP, remained enlarged, indicating that the essentiality of c-di-AMP is bypassed in this strain potentially through a different mechanism. Here, we show that AlsT is an efficient glutamine transporter in *S. aureus* (Fig. 2). These findings indicate that eliminating or reducing the ability of *S. aureus* to take up glutamine from rich growth medium rescues the growth of an *S. aureus* unable to produce c-di-AMP. There are several (not mutually exclusive) possibilities how preventing glutamine uptake could rescue the growth of a c-di-AMP null strain in rich medium. Glutamine as well as proline have been shown to accumulate in *S. aureus* under NaCl stress conditions (Anderson & Witter, 1982). While it has been suggested that the glutamine accumulation is due to synthesis rather than uptake (Anderson & Witter, 1982), these data highlight that glutamine likely plays an important role in osmotic regulation in *S. aureus.* Despite the cell size not being restored in the *dacA/alsT* suppressor strain, blocking glutamine uptake could potentially still help bacteria to better balance their cellular osmolality during growth in rich medium in the absence of c-di-AMP. Another possible explanation how eliminating glutamine uptake could allow *S. aureus* to grow in the absence of c-di-AMP could be related to changes in metabolism and TCA cycle activity. In *L. monocytogenes,* an increased flux of pyruvate into the TCA cycle has been described for a strain unable to produce c-di-AMP (Sureka *et al.*, 2014). As a consequence, an accumulation of citrate and increased carbon flux into glutamine/glutamate was observed, which resulted in a metabolic imbalance and growth defect (Sureka *et al.*, 2014). Perhaps similar to the observations in *L. monocytogenes*, the absence of c-di-AMP could also boost TCA cycle activity in *S. aureus*, thus leading to glutamine accumulation and a metabolic imbalance. Hence, the lack of c-di-AMP combined with active glutamine uptake could fuel the bacterial metabolism and the resulting metabolic imbalance might become toxic to the cell, similar as observed for *L. monocytogenes* (Sureka *et al.*, 2014, Whiteley *et al.*, 2017).

In a recent study investigating genetic determinants required for eDNA release during biofilm formation, it was found that inactivation of GdpP as well as of AlsT, resulted in a significant decrease in eDNA release and in an increase in resistance to Congo red (DeFrancesco *et al.*, 2017). Therefore, inactivation of AlsT and preventing/reducing glutamine uptake might lead to alterations in the bacterial cell wall that make bacteria more resistant to cell lysis. Such changes could also be an advantage during osmotic stress or c-di-AMP deficiency. Indeed, we have recently shown a correlation between specific changes in the peptidoglycan structure and the NaCl stress resistance in *S. aureus* (Schuster *et al.*, 2019). In addition, since the cellular c-di-AMP levels are significantly higher in the *gdpP* as well as the *alsT* mutant strains compared to WT (Fig. 7B and Fig. 8A), the underlying mechanistic bases for the decrease in eDNA release observed for the *gdpP* and *alsT* mutant strains might be related.

The actual stimuli and underlying molecular mechanisms that regulate c-di-AMP production in bacterial cells are at the moment poorly understood. As part of this study, we show that ammonium and AlsT-mediated glutamine uptake but not GltS-mediated glutamate uptake negatively impacts c-di-AMP production (Fig. 7). Changes in cellular c-di-AMP levels depending on the presence of ammonium, glutamine or glutamate have already been reported for *B. subtilis* (Gundlach *et al.*, 2015). For *B. subtilis* it has been suggested that glutamine and to some extent ammonium uptake stimulates c-di-AMP production (Gundlach *et al.*, 2015). Here we show that in *S. aureus* ammonium and glutamine uptake leads to an inhibition of c-di-AMP production rather than glutamate promoting its synthesis (Fig. 7). The decrease in c-di-AMP production in the presence of ammonium or glutamine is likely achieved by reducing the activity of the c-di-AMP cyclase DacA and not by activation of the c-di-AMP specific phosphodiesterase GdpP. This conclusion is based on our observation that the cellular c-di-AMP levels also decreased in a *gdpP* mutant strain upon addition of glutamine or ammonium (Fig. 8A). Current evidence suggests that the activity of DacA can be regulated through the interaction with two proteins: the membrane anchored and proposed DacA regulator protein YbbR (also name CdaR in other bacteria) and the phosphoglucomutase enzyme GlmM (Tosi *et al.*, 2019, Zhu *et al.*, 2016, Gundlach *et al.*, 2015, Pham *et al.*, 2016). We could exclude that the observed reduction in cellular c-di-AMP levels in the presence of ammonium or glutamine is mediated by YbbR, as <u>a</u> *ybbR* mutant showed a similar decrease in the c-di-AMP levels as observed for the WT strain (Fig. 8B). GlmM has been shown to be a negative regulator of DacA activity both *in vivo* and *in vitro* (Tosi *et al.*, 2019, Pham *et al.*, 2018). However, since GlmM is likely an essential enzyme in *S. aureus*, we were unable to construct a *glmM* mutant and test its involvement in the observed repression of c-di-AMP synthesis in the presence of ammonium or glutamine as we did for the *gdpP* and *ybbR* mutant strains. Nevertheless, with this work, we not only identified main glutamine and glutamate transporters in *S. aureus,* but we also linked the c-di-AMP signalling network to central nitrogen metabolism in *S. aureus.* It will be interesting to determine in future studies the mechanistic bases for the observed changes in cellular c-di-AMP levels depending on ammonium and glutamine uptake and the involvement of GlmM or other factors in this process.

## Experimental Procedures

### Bacterial strains and culture conditions

Bacterial strains used in this study are listed in Table 1. *S. aureus* strains were grown in Tryptic Soy Broth (TSB), Tryptic Soy Agar (TSA) or Glucose Defined Medium (GDM). GDM was prepared similar to the chemically defined medium (CDM) reported in an earlier study (Zeden *et al.*, 2018), with some modifications. The detailed content of the GDM (which contains essential vitamins, trace metals, amino acids and glucose as a carbon source but lacks ammonium, glutamine or glutamate as a potential nitrogen or amino acid source) is shown in Table S1. In addition to the GDM, GDM containing glutamine (GDM+Gln), glutamate (GDM+Glu), ammonium (GDM+NH_3_), glutamine and ammonium (GDM+Gln+NH_3_) or glutamate and ammonium (GDM+Glu+NH_3_) were used as part of this study (for exact composition see Table S1). Where indicated, certain amino acids were removed from the GDM recipe during uptake assays and when needed the TSB was supplemented with 0.4 M NaCl. *Escherichia coli* strains were grown in Lysogeny Broth (LB). Where appropriate, antibiotics and/or inducers were added to the media at the following concentration: 200 ng/ml anhydrotetracycline (Atet), 90 μg/ml Kanamycin (Kan), 10 μg/ml Erythromycin (Erm), 7.5 or 10 μg/ml Chloramphenicol (Cam), Ampicillin (Amp) 100 μg/ml.

**Table 1:**
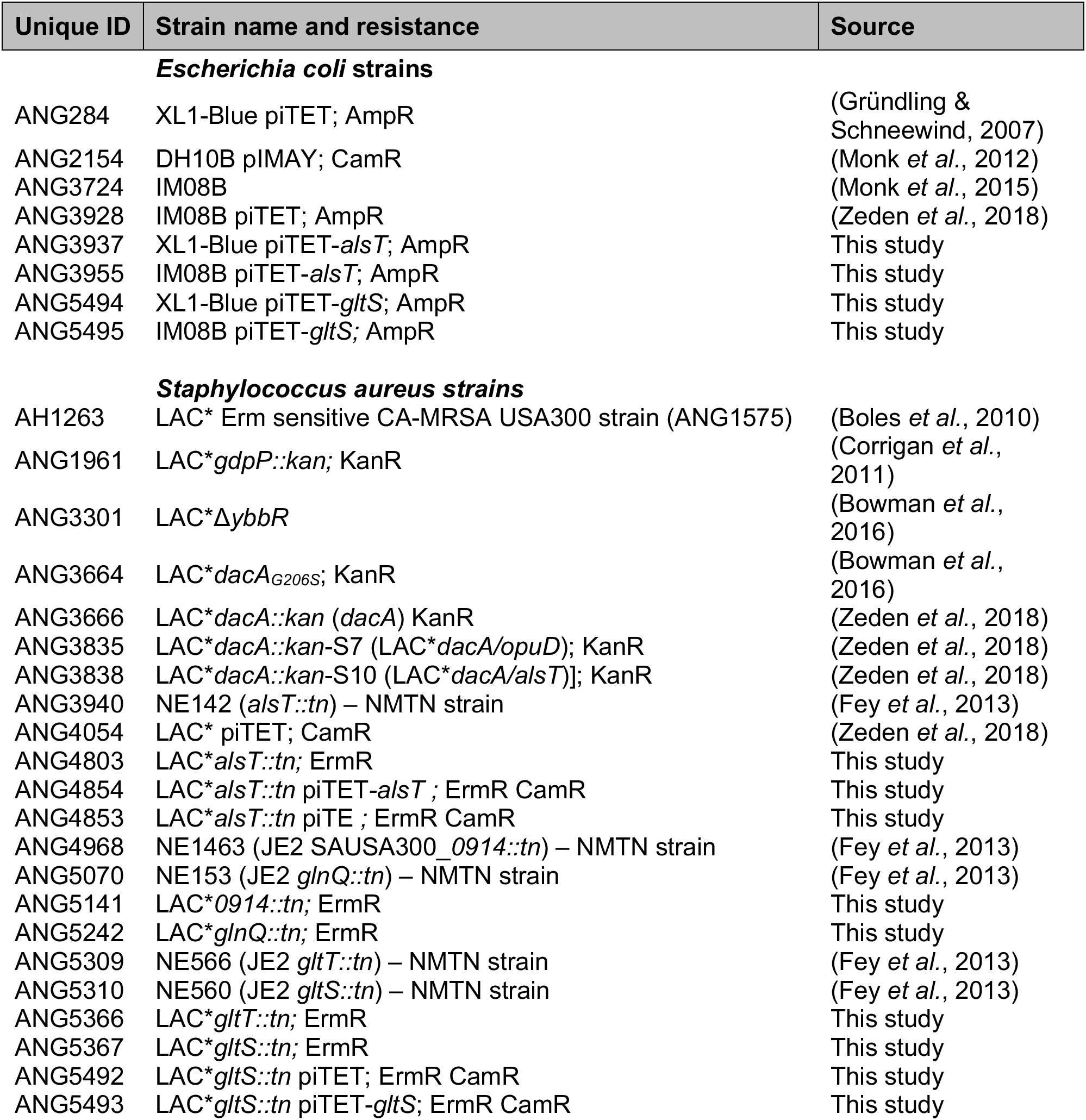
Bacterial strains used in this study

| Unique ID | Strain name and resistance | Source |
| --- | --- | --- |
| ***Escherichia coli* strains** |  |  |
| ANG284 | XL1-Blue pTET; AmpR | (Gründling & Schneewind, 2007) |
| ANG2154 | DH10B pIMAY; CamR | (Monk *et al.*, 2012) |
| ANG3724 | IM08B | (Monk *et al.*, 2015) |
| ANG3928 | IM08B pTET; AmpR | (Zeden *et al.*, 2018) |
| ANG3937 | XL1-Blue pTET-*alsT*; AmpR | This study |
| ANG3955 | IM08B pTET-*alsT*; AmpR | This study |
| ANG5494 | XL1-Blue pTET-*gltS*; AmpR | This study |
| ANG5495 | IM08B pTET-*gltS*; AmpR | This study |
| ***Staphylococcus aureus* strains** |  |  |
| AH1263 | LAC* Erm sensitive CA-MRSA USA300 strain (ANG1575) | (Boles *et al.*, 2010) |
| ANG1961 | LAC* *gdpP::kan*; KanR | (Corrigan *et al.*, 2011) |
| ANG3301 | LAC*Δ*ybbR* | (Bowman *et al.*, 2016) |
| ANG3664 | LAC* *dacA* <sub>G206S</sub>; KanR | (Bowman *et al.*, 2016) |
| ANG3666 | LAC* *dacA::kan* (*dacA*) KanR | (Zeden *et al.*, 2018) |
| ANG3835 | LAC* *dacA::kan-S7* (LAC* *dacA/opuD*); KanR | (Zeden *et al.*, 2018) |
| ANG3838 | LAC* *dacA::kan-S10* (LAC* *dacA/alsT*); KanR | (Zeden *et al.*, 2018) |
| ANG3940 | NE142 (*alsT::tn*) – NMTN strain | (Fey *et al.*, 2013) |
| ANG4054 | LAC* pTET; CamR | (Zeden *et al.*, 2018) |
| ANG4803 | LAC* *alsT::tn*; ErmR | This study |
| ANG4854 | LAC* *alsT::tn* pTET- *alsT*; ErmR CamR | This study |
| ANG4853 | LAC* *alsT::tn* pTE; ErmR CamR | This study |
| ANG4968 | NE1463 (JE2 SAUSA300_0914::tn) – NMTN strain | (Fey *et al.*, 2013) |
| ANG5070 | NE153 (JE2 *glnQ::tn*) – NMTN strain | (Fey *et al.*, 2013) |
| ANG5141 | LAC*0914::tn; ErmR | This study |
| ANG5242 | LAC* *glnQ::tn*; ErmR | This study |
| ANG5309 | NE566 (JE2 *gltT::tn*) – NMTN strain | (Fey *et al.*, 2013) |
| ANG5310 | NE560 (JE2 *gltS::tn*) – NMTN strain | (Fey *et al.*, 2013) |
| ANG5366 | LAC* *gltT::tn*; ErmR | This study |
| ANG5367 | LAC* *gltS::tn*; ErmR | This study |
| ANG5492 | LAC* *gltS::tn* pTET; ErmR CamR | This study |
| ANG5493 | LAC* *gltS::tn* pTET- *gltS*; ErmR CamR | This study |

### Bacterial strain construction

All strains used in this study are listed in Table 1 and primers used in this study are listed in Table 2. The transposon insertion sites in the Nebraska transposon mutant library (NTML) strains (Fey *et al.*, 2013) used as part of this study were confirmed by PCR and sequencing. The transposon and surrounding regions were moved by phage transduction using phage 85 into the *S. aureus* LAC* strain background. This resulted in the generation of *S. aureus* strains LAC\**alsT∷tn* (*SAUSA300_1252∷tn*; ANG4803), LAC\**0914∷tn* (*SAUSA300_0914∷tn*; ANG5141), LAC\**glnQ∷tn* (*SAUSA300_1807∷tn;* ANG5070), LAC\**gltT∷tn* (*SAUSA300_2329∷tn*; ANG5366) and LAC\**gltS∷tn* (*SAUSA300_2291∷tn*; ANG5367). The transposon insertion in the respective gene was again confirmed by PCR and sequencing. For complementation analysis, the Atet inducible single copy integration plasmids piTET-*alsT* and piTET-*gltS* were constructed. To this end, *alsT* (*SAUSA300_1252*) and *gltS* (*SAUSA300_2291*) were amplified using LAC* chromosomal DNA and primers ANG2250/ANG2251 and ANG3209/ANG3210, respectively. The products as well as piTET were digested with AvrII and SacII and then ligated. Plasmid piTET-*alsT* was recovered in *E. coli* strain XL1-Blue (yielding strain ANG3937), shuttled through *E. coli* strain IM08B (strain ANG3955) and then introduced into LAC\**alsT∷tn* (ANG4803), yielding strain LAC\**alsT∷tn* piTET*-alsT* (ANG4854). As a control, plasmid piTET was also introduced into LAC\**alsT∷tn* (ANG4803) yielding strain LAC\**alsT∷tn* piTET (ANG4853). Plasmid piTET-*gltS* was transformed into *E. coli* XL1-Blue (yielding strain ANG5494), shuttled through *E. coli* IM08B (yielding strain ANG5495) and transformed into *LAC*gltS∷tn*, yielding the complement strain LAC\**gltS∷tn* piTET-*gltS* (ANG5493). As a control, the piTET plasmid was transformed into LAC\**gltS∷tn* strain, yielding the strain LAC\**gltS*:tn piTET (ANG5492). Correct plasmid integration into the *geh* locus was confirmed by PCR and the sequences of all plasmid inserts were confirmed by fluorescent automated sequencing.

**Table 2:**
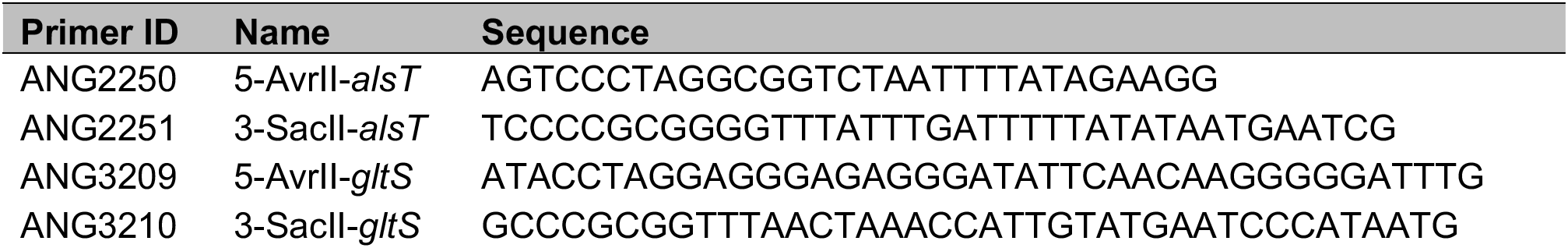
Cloning primers used in this study

### Bacterial growth curves and amino acid analysis in culture supernatants

*S. aureus* strains LAC* and LAC\**alsT∷tn* were grown overnight in TSB supplemented with 10 μg/ml erythromycin where appropriate. Overnight cultures were then diluted to an OD_600_ of 0.01 into 50 ml of fresh TSB. Cultures were incubated at 37°C with aeration, and OD_600_ values determined every hour. The experiment was performed with three biological replicates and the average OD_600_ values and standard deviations (SDs) were plotted. Using the same cultures, supernatant samples were prepared at the 0, 6, 10 and 12 h time points and amino acid levels determined as previously described using an amino acid analyser (Halsey *et al.*, 2017). For measuring the growth of *S. aureus* strains LAC*, LAC* piTET, LAC\**alsT∷tn*, LAC\**alsT∷tn* piTET, LAC\**alsT∷tn* piTET-*alsT* and LAC\**gltS∷tn* in GDM, GDM+Gln, GDM+Glu, GDM+NH_3_, GDM+Gln+NH_3_ or GDM+Glu+NH_3_, the bacteria were grown overnight in TSB medium supplemented with chloramphenicol and erythromycin where appropriate. Next day, bacteria from a 1 ml aliquot were washed twice with PBS and diluted to OD_600_ of 0.005 in the indicated GDM. LAC* WT, LAC\**alsT∷tn* and LAC\**gltS∷tn* were grown in GDM, GDM+Gln, GDM+Glu, GDM+NH_3_, GDM+Gln+NH_3_ and GDM+Glu+NH_3_ while LAC* piTET, LAC\**alsT∷tn* piTET and LAC\**alsT∷tn* piTET-*alsT* were grown in GDM+Gln supplemented with 200 ng/ml Atet. One hundred μl of the diluted cultures (six technical replicates) were transferred into wells of a 96-well plate and the plate was then incubated with shaking (500 rpm) in a plate reader and OD_600_ readings determined every 30 min. The average values of the technical replicates were determined for each strain. The experiment was performed three times and the average readings and standard deviations were plotted.

### γ-L-glutamyl hydrazide susceptibility assay

The susceptibility of *S. aureus* LAC*, LAC\**alsT∷tn* (ANG4803), LAC\**0914∷tn* (ANG5141) and LAC\**glnQ∷tn* (ANG5242) to the toxic glutamine analogue **γ**-L-glutamyl hydrazide (Alfa Aesar, MA, USA) was determined using a similar method as previously reported (Zhu *et al.*, 2009). Briefly, the different strains were grown overnight at 37°C in 5 ml TSB medium, supplemented with 10 μg/ml erythromycin where appropriate. Next day, the bacteria were washed twice with PBS, diluted to an OD_600_ of 0.005 in GDM+NH_3_. Next, 10 μl of water (0 mM control) or 10 μl of a γ-L-glutamyl hydrazide solution dissolved in water was added to 0.99 ml aliquots of these bacterial suspension to give a final concentration of 20, 40, 60 or 80 μg/ml, respectively. One hundred μl were subsequently transferred in four replicates into wells of a 96 well plates and the plate incubated at 37°C with shaking (500 rpm) in a plate reader and OD_600_ readings determined every 10 min for 12 h. The experiment was performed three times and the average OD_600_ values of the three experiments presented as growth curves. The average values and SDs of the OD_600_ values from the 7 h time point were also plotted against the different **γ**-L-glutamyl hydrazide concentrations.

### Microscopic analysis and cell size measurements

The microscopic analysis to determine bacterial cell sizes was performed essentially as previously described (Zeden *et al.*, 2018). Briefly, *S. aureus* strains LAC*, LAC\**dacA∷kan*, LAC\**dacA*G206S, LAC\**dacA/opuD* (ANG3835) and LAC\**dacA/alsT* (ANG3838) were grown overnight at 37°C in TSB or TSB supplemented with 0.4 M NaCl where stated. Next day, the cultures were diluted to an OD_600_ of 0.01 and grown for 3 h at 37°C to mid-exponential phase (OD_600_ of 0.5-0.9). One hundred μl of these cultures were then stained for 20 min at 37°C with Vancomycin-BODIPY FL at a final concentration of 2 μg/ml. One and a half μl of each sample was spotted onto a thin 1.5% agarose gel patch prepared in H_2_O or in 0.4 M NaCl and the bacteria subsequently imaged at 1000 × magnification using an Axio Imager A2 Zeiss microscope equipped with a GFP filter set. Images were acquired using the ZEN 2012 (blue edition) software. The bacterial cell diameters were determined using the Fiji software. Only non-dividing cells (cells without any obvious fluorescent dots or lines at the mid-cell), were used for cell diameter measurements. The cell diameters of 50 cells were measured and the average cell diameter determined. The experiment was conducted three or four times (as indicated in the figure legend) and the averages and standard deviations of the average cell diameters plotted.

### Uptake assays using ^14^C-labelled amino acids

Uptake assays were conducted as previously described with some minor modifications (Zeden *et al.*, 2018). Briefly, *S. aureus* strains were streaked on TSA or TSA 0. 4M NaCl plates with appropriate antibiotics and the plates incubated overnight at 37°C. Bacteria were subsequently scraped off from the plates and suspended in 1 ml PBS pH 7.4 buffer and the OD_600_ determined. Fifty ml of GDM+Glu+NH_3_ (where indicated with 200 ng/ml of the inducer Atet added) was inoculated with the appropriate bacterial suspensions to an OD_600_ of 0.05. The cultures were grown at 37°C to an OD_600_ between 0.4 and 0.9 and bacteria from an OD_600_ equivalent of 8 were harvested by centrifugation for 10 min at 19,000 × g at RT. Supernatants were discarded and the bacterial pellets were suspended in 2 ml of GDM+NH_3_ (for glutamine and glutamate uptake assays) or GDM+Glu+NH_3_ without serine (for serine uptake assays). The OD_600_ of the cell suspensions were measured and the cells diluted to an OD_600_ of approximately 1. The OD_600_ was re-measured and this measurement used for normalization purposes. Five hundred and fifty μl of these cell suspensions were aliquoted into 50 ml conical tubes and 100 μl used to measure the background radiation, by filtering the cells onto a nitrocellulose membrane filter, followed by a wash step with 16 ml PBS. Then, 6.2 μl of glutamine, L-[14C(U)] (Hartmann Analytic, MC1124), glutamic acid, L-[14C(U)] (Hartmann Analytic, MC156), or serine L-[14C(U)] (Hartmann Analytic, MC265) was added to the remaining 450 μl sample. One hundred μl aliquots were filtered 0, 3, 6 and 9 min after addition of the radiolabelled amino acid and the filters were then washed two times with 16 ml of PBS pH 7.4. The filters were subsequently dissolved in 9 ml of scintillation cocktail Filter Count (Perkin Elmer) and the radioactivity measured in counts per minute (CPM) using a Wallac 1409 DSA liquid scintillation counter. The CPMs were then normalized to the OD_600_ reading of the final cell suspension and the means and standard deviations of the CPM/ml OD_600_ = 1 of three or four (as indicated in the figure legends) independent experiments were plotted.

### Determination of cellular c-di-AMP levels by competitive ELISA

Intracellular c-di-AMP levels in WT LAC* and the indicated *S. aureus* mutant strains were determined using a previously described competitive ELISA method (Underwood *et al.*, 2014) and a slightly modified method for the preparation of *S. aureus* samples (Bowman *et al.*, 2016). Briefly, single colonies of the WT LAC* strain were picked from TSA plates and used to inoculate 5 ml of GDM, GDM+Gln, GDM+Glu, GDM+NH_3_, GDM+Gln+NH_3_ and GDM+Glu+NH_3_. Colonies of the strains LAC\**gdpP∷kan* and LAC*Δ*ybbR* were inoculated into 5 ml of GDM, GDM+Glu and GDM+Gln. Colonies of strains LAC* piTET, LAC\**alsT∷tn* piTET and LAC\**alsT∷tn* piTET-*alsT* were inoculated into GDM+Gln containing 200 ng/ml Atet and colonies of strains LAC* piTET, LAC\**gltS∷tn* piTET and LAC\**gltS∷tn* piTET-*gltS* were inoculated into GDM+Glu supplemented with 200 ng/ml Atet. All cultures were incubated for 18 h at 37°C with shaking. Next, bacteria from 4.5 ml culture were collected by centrifugation, washed three times with PBS and subsequently suspended in 0.75 to 1 ml 50 mM Tris pH 8 buffer supplemented with 20 ng/ml lysostaphin and the cells were lysed by bead beating. The lysates were cleared by centrifugation for 5 min at 17,000 × g and the supernatant transferred to a new tube. A small sample aliquot was removed, and the protein concentration determined for normalization purposes using a Pierce BCA protein assay kit (Thermo Scientific, Waltham, MA, USA). The remainder of the sample was heated to 95°C for 10 min. For the competitive ELISA assay, the samples were diluted to a protein concentration of 100, 200, 400 or 500 μg/ml as, appropriate. ELISA plates were prepared by adding 100 μl of coating buffer (50 mM Na_2_C0_3_, 50 mM NaHCO_3_, pH 9.6) containing 10 μg/ml of the c-di-AMP binding protein CpaA_SP_ to each well of a 96 well NUNC MaxiSorp plate (Thermo Scientific, Waltham, MA, USA) and the plate was incubated for approximately 18 h at 4°C. Next, the plate was washed three times with 200 μl PBST pH 7.4 (10 mM Na_2_HPO_4_, 1.8 mM KH_2_PO_4_ 137 mM NaCl, 2.7 mM KCl, 0.05% (v/v) Tween 20), blocked for 1 h at 18°C with 150 μl blocking solution (1% BSA in PBS pH 7.4) and washed three times with 200 μl PBST. Fifty μl of the samples (three biological replicates and three technical replicates) or standards (two technical replicates) were mixed with 50 μl of a 50 nM biotinylated c-di-AMP solution prepared in 50 mM Tris pH 8 buffer. For the standard curve, c-di-AMP standards were prepared in 50 mM Tris pH 8 buffer at concentrations of 0, 12.5, 25, 37.5, 50, 75, 100, 200 nM. Following the addition of the samples and the standards, the plate was incubated for 2 h at 18°C and then washed three times with PBST. Next, 100 μl of a high sensitivity streptavidin-HRP solution (Thermo Scientific, Waltham, MA, USA) diluted 1:500 in PBS was added to each well and the plate was incubated for 1 h at 18°C. The plate was washed again 3 × with 200 μl PBST and 100 μl of a developing solution (0.103 M NaHPO_4_, 0.0485 M citric acid, 500 mg/l o-phenylenediamine dihydrochloride, 0.03% H_2_O_2_) was added to each well and the plate incubated for 15 min at 18°C. The reaction was then stopped by adding 100 μl of 2 M H_2_SO_4_ solution. The absorbance was measured in a plate reader at a wavelength of 490 nm and c-di-AMP concentrations were calculated as ng c-di-AMP / mg protein.

## Acknowledgments

This work was funded by the Wellcome Trust grants 100289/Z/12/Z and 210671/Z/18/Z to AG and the NIH/NIAID grants P01AI083211 and R01AI125588 to PDF and VCT, respectively. MSZ was supported by a Medical Research Council Centre for Molecular Bacteriology and Infection (MRC CMBI) studentship. The funders had no role in the study design, data collection and interpretation, or the decision to submit the work for publication. The authors have no conflicts of interest to declare. The data supporting the findings of this study will be openly available.

## Author contribution

MSZ, IK and AG designed the study, MSZ, IK and CFS acquired the data, MSZ, IK, CFS, VCT, PDF and AG, designed experiments, analyzed and interpreted the data, MSZ, IK and AG prepared the figures and wrote the original draft of the manuscript. All authors approved the final version of the manuscript.

## Abbreviated Summary

A large number of amino acid transporters and oligopeptide permeases are encoded in bacterial genomes. However, their actual substrate specificity and functions are hard to predict bioinformatically. In this study, we report that GltS and AlsT are main glutamate and glutamine transporters in *Staphylococcus aureus*, respectively and show that glutamine and ammonium uptake inhibits the production of the nucleotide signalling molecule c-di-AMP.

## Notes

#### Summary of Updates

This revised version contains additional experiment. Growth of WT and mutant S. aureus strains in GDM with glutamine/glutamate/ammonium as well as additional c-di-AMP level measurements.

## References

Anderson, C.B., and Witter, L.D. (1982) Glutamine and proline accumulation by *Staphylococcus aureus* with reduction in water activity. Appl Environ Microbiol 43: 1501–1503.

Bai, Y., Yang, J., Eisele, L.E., Underwood, A.J., Koestler, B.J., Waters, C.M., Metzger, D.W., and Bai, G. (2013) Two DHH subfamily 1 proteins in *Streptococcus pneumoniae* possess cyclic di-AMP phosphodiesterase activity and affect bacterial growth and virulence. J Bacteriol 195: 5123–5132.

Bai, Y., Yang, J., Zarrella, T.M., Zhang, Y., Metzger, D.W., and Bai, G. (2014) Cyclic di-AMP impairs potassium uptake mediated by a cyclic di-AMP binding protein in *Streptococcus pneumoniae*. J Bacteriol 196: 614–623.

Boles, B.R., Thoendel, M., Roth, A.J., and Horswill, A.R. (2010) Identification of genes involved in polysaccharide-independent *Staphylococcus aureus* biofilm formation. PLoS One 5: e10146.

Bowman, L., Zeden, M.S., Schuster, C.F., Kaever, V., and Gründling, A. (2016) New Insights into the Cyclic Di-adenosine Monophosphate (c-di-AMP) Degradation Pathway and the Requirement of the Cyclic Dinucleotide for Acid Stress Resistance in *Staphylococcus aureus*. J Biol Chem 291: 26970–26986.

Chin, K.H., Liang, J.M., Yang, J.G., Shih, M.S., Tu, Z.L., Wang, Y.C., Sun, X.H., Hu, N.J., Liang, Z.X., Dow, J.M., Ryan, R.P., and Chou, S.H. (2015) Structural Insights into the Distinct Binding Mode of Cyclic Di-AMP with SaCpaA_RCK. Biochemistry 54: 4936–4951.

Commichau, F.M., Heidemann, J.L., Ficner, R., and Stülke, J. (2019) Making and Breaking of an Essential Poison: the Cyclases and Phosphodiesterases That Produce and Degrade the Essential Second Messenger Cyclic di-AMP in Bacteria. Journal of Bacteriology 201: e00462–00418.

Corrigan, R.M., Abbott, J.C., Burhenne, H., Kaever, V., and Gründling, A. (2011) c-di-AMP is a new second messenger in *Staphylococcus aureus* with a role in controlling cell size and envelope stress. PLoS Pathog 7: e1002217.

Corrigan, R.M., Bowman, L., Willis, A.R., Kaever, V., and Gründling, A. (2015) Cross-talk between two nucleotide-signaling pathways in *Staphylococcus aureus*. J Biol Chem 290: 5826–5839.

Corrigan, R.M., Campeotto, I., Jeganathan, T., Roelofs, K.G., Lee, V.T., and Gründling, A. (2013) Systematic identification of conserved bacterial c-di-AMP receptor proteins. Proc Natl Acad Sci U S A 110: 9084–9089.

Crooke, A.K., Fuller, J.R., Obrist, M.W., Tomkovich, S.E., Vitko, N.P., and Richardson, A.R. (2013) CcpA-independent glucose regulation of lactate dehydrogenase 1 in *Staphylococcus aureus*. PLoS One 8: e54293.

DeFrancesco, A.S., Masloboeva, N., Syed, A.K., DeLoughery, A., Bradshaw, N., Li, G.W., Gilmore, M.S., Walker, S., and Losick, R. (2017) Genome-wide screen for genes involved in eDNA release during biofilm formation by *Staphylococcus aureus*. Proc Natl Acad Sci U S A 114: E5969–E5978.

Devaux, L., Sleiman, D., Mazzuoli, M.V., Gominet, M., Lanotte, P., Trieu-Cuot, P., Kaminski, P.A., and Firon, A. (2018) Cyclic di-AMP regulation of osmotic homeostasis is essential in Group B *Streptococcus*. PLoS Genet 14: e1007342.

Fahmi, T., Faozia, S., Port, G., and Cho, K.H. (2019) The Second Messenger c-di-AMP Regulates Diverse Cellular Pathways Involved in Stress Response, Biofilm Formation, Cell Wall Homeostasis, SpeB Expression and Virulence in *Streptococcus pyogenes*. Infect Immun 87: e00147–00119.

Fey, P.D., Endres, J.L., Yajjala, V.K., Widhelm, T.J., Boissy, R.J., Bose, J.L., and Bayles, K.W. (2013) A genetic resource for rapid and comprehensive phenotype screening of nonessential *Staphylococcus aureus* genes. MBio 4: e00537–00512.

Fridkin, S.K., Hageman, J.C., Morrison, M., Sanza, L.T., Como-Sabetti, K., Jernigan, J.A., Harriman, K., Harrison, L.H., Lynfield, R., Farley, M.M., and Active Bacterial Core Surveillance Program of the Emerging Infections Program, N. (2005) Methicillin-resistant *Staphylococcus aureus* disease in three communities. N Engl J Med 352: 1436–1444.

Fuller, J.R., Vitko, N.P., Perkowski, E.F., Scott, E., Khatri, D., Spontak, J.S., Thurlow, L.R., and Richardson, A.R. (2011) Identification of a lactate-quinone oxidoreductase in *Staphylococcus aureus* that is essential for virulence. Front Cell Infect Microbiol 1: 19.

Gründling, A., and Schneewind, O. (2007) Genes required for glycolipid synthesis and lipoteichoic acid anchoring in *Staphylococcus aureus*. J Bacteriol 189: 2521–2530.

Gundlach, J., Commichau, F.M., and Stülke, J. (2018) Perspective of ions and messengers: an intricate link between potassium, glutamate, and cyclic di-AMP. Curr Genet 64: 191–195.

Gundlach, J., Herzberg, C., Hertel, D., Thurmer, A., Daniel, R., Link, H., and Stülke, J. (2017a) Adaptation of *Bacillus subtilis* to Life at Extreme Potassium Limitation. MBio 8: e00861–00817.

Gundlach, J., Herzberg, C., Kaever, V., Gunka, K., Hoffmann, T., Weiss, M., Gibhardt, J., Thurmer, A., Hertel, D., Daniel, R., Bremer, E., Commichau, F.M., and Stülke, J. (2017b) Control of potassium homeostasis is an essential function of the second messenger cyclic di-AMP in *Bacillus subtilis*. Sci Signal 10: eaal3011.

Gundlach, J., Krüger, L., Herzberg, C., Turdiev, A., Poehlein, A., Tascon, I., Weiß, M., Hertel, D., Daniel, R., Hänelt, I., Lee, V.T., and Stülke, J. (2019) Sustained sensing in potassium homeostasis: Cyclic di-AMP controls potassium uptake by KimA at the levels of expression and activity. J Biol Chem 294: 9605–9614.

Gundlach, J., Mehne, F.M., Herzberg, C., Kampf, J., Valerius, O., Kaever, V., and Stülke, J. (2015) An Essential Poison: Synthesis and Degradation of Cyclic Di-AMP in *Bacillus subtilis*. J Bacteriol 197: 3265–3274.

Halsey, C.R., Lei, S., Wax, J.K., Lehman, M.K., Nuxoll, A.S., Steinke, L., Sadykov, M., Powers, R., and Fey, P.D. (2017) Amino Acid Catabolism in *Staphylococcus aureus* and the Function of Carbon Catabolite Repression. MBio 8: e01434–01416.

Hengge, R. (2009) Principles of c-di-GMP signalling in bacteria. Nat Rev Microbiol 7: 263–273.

Huynh, T.N., Choi, P.H., Sureka, K., Ledvina, H.E., Campillo, J., Tong, L., and Woodward, J.J. (2016) Cyclic di-AMP targets the cystathionine beta-synthase domain of the osmolyte transporter OpuC. Mol Microbiol 102: 233–243.

Kelly, B., and O’Neill, L.A. (2015) Metabolic reprogramming in macrophages and dendritic cells in innate immunity. Cell Res 25: 771–784.

Kim, H., Youn, S.J., Kim, S.O., Ko, J., Lee, J.O., and Choi, B.S. (2015) Structural Studies of Potassium Transport Protein KtrA Regulator of Conductance of K+ (RCK) C Domain in Complex with Cyclic Diadenosine Monophosphate (c-di-AMP). J Biol Chem 290: 16393–16402.

Kluytmans, J., van Belkum, A., and Verbrugh, H. (1997) Nasal carriage of *Staphylococcus aureus*: epidemiology, underlying mechanisms, and associated risks. Clin Microbiol Rev 10: 505–520.

Lehman, M.K., Nuxoll, A.S., Yamada, K.J., Kielian, T., Carson, S.D., and Fey, P.D. (2019) Protease-Mediated Growth of *Staphylococcus aureus* on Host Proteins Is *opp3* Dependent. MBio 10: e02553–02518.

Mehne, F.M., Gunka, K., Eilers, H., Herzberg, C., Kaever, V., and Stülke, J. (2013) Cyclic di-AMP homeostasis in *Bacillus subtilis*: both lack and high level accumulation of the nucleotide are detrimental for cell growth. J Biol Chem 288: 2004–2017.

Monk, I.R., Shah, I.M., Xu, M., Tan, M.W., and Foster, T.J. (2012) Transforming the untransformable: application of direct transformation to manipulate genetically *Staphylococcus aureus* and *Staphylococcus epidermidis*. MBio 3: e00277–00211.

Monk, I.R., Tree, J.J., Howden, B.P., Stinear, T.P., and Foster, T.J. (2015) Complete Bypass of Restriction Systems for Major *Staphylococcus aureus* Lineages. MBio 6: e00308–00315.

Moscoso, J.A., Schramke, H., Zhang, Y., Tosi, T., Dehbi, A., Jung, K., and Gründling, A. (2015) Binding of Cyclic Di-AMP to the *Staphylococcus aureus* Sensor Kinase KdpD Occurs via the Universal Stress Protein Domain and Downregulates the Expression of the Kdp Potassium Transporter. J Bacteriol 198: 98–110.

Pham, H.T., Nhiep, N.T.H., Vu, T.N.M., Huynh, T.N., Zhu, Y., Huynh, A.L.D., Chakrabortti, A., Marcellin, E., Lo, R., Howard, C.B., Bansal, N., Woodward, J.J., Liang, Z.X., and Turner, M.S. (2018) Enhanced uptake of potassium or glycine betaine or export of cyclic-di-AMP restores osmoresistance in a high cyclic-di-AMP *Lactococcus lactis* mutant. PLoS Genet 14: e1007574.

Pham, H.T., and Turner, M.S. (2019) Onwards and [K(+)]upwards: a new potassium importer under the spell of cyclic-di-AMP. J Bacteriol 201: e00150–00119.

Pham, T.H., Liang, Z.X., Marcellin, E., and Turner, M.S. (2016) Replenishing the cyclic-di-AMP pool: regulation of diadenylate cyclase activity in bacteria. Curr Genet 62: 731–738.

Quintana, I.M., Gibhardt, J., Turdiev, A., Hammer, E., Commichau, F.M., Lee, V.T., Magni, C., and Stülke, J. (2019) The KupA and KupB proteins of *Lactococcus lactis* IL1403 are novel c-di-AMP receptor proteins responsible for potassium uptake. J Bacteriol 201: e00028–00019.

Richardson, A.R., Libby, S.J., and Fang, F.C. (2008) A nitric oxide-inducible lactate dehydrogenase enables *Staphylococcus aureus* to resist innate immunity. Science 319: 1672–1676.

Rocha, R., Teixeira-Duarte, C.M., Jorge, J.M.P., and Morais-Cabral, J.H. (2019) Characterization of the molecular properties of KtrC, a second RCK domain that regulates a Ktr channel in *Bacillus subtilis*. J Struct Biol 205: 34–43.

Römling, U. (2008) Great times for small molecules: c-di-AMP, a second messenger candidate in Bacteria and Archaea. Sci Signal 1: pe39.

Schuster, C.F., Bellows, L.E., Tosi, T., Campeotto, I., Corrigan, R.M., Freemont, P., and Gründling, A. (2016) The second messenger c-di-AMP inhibits the osmolyte uptake system OpuC in *Staphylococcus aureus*. Sci Signal 9: ra81.

Schuster, C.F., Wiedemann, D.M., Kirsebom, F.C.M., Santiago, M., Walker, S., and Gründling, A. (2019) High-throughput transposon sequencing highlights the cell wall as an important barrier for osmotic stress in methicillin resistant *Staphylococcus aureus* and underlines a tailored response to different osmotic stressors. Mol Microbiol: DOI: 10.1111/mmi.14433.

Schuurman-Wolters, G.K., and Poolman, B. (2005) Substrate specificity and ionic regulation of GlnPQ from *Lactococcus lactis*. An ATP-binding cassette transporter with four extracytoplasmic substrate-binding domains. J Biol Chem 280: 23785–23790.

Spahich, N.A., Vitko, N.P., Thurlow, L.R., Temple, B., and Richardson, A.R. (2016) *Staphylococcus aureus* lactate-and malate-quinone oxidoreductases contribute to nitric oxide resistance and virulence. Mol Microbiol 100: 759–773.

Sureka, K., Choi, P.H., Precit, M., Delince, M., Pensinger, D., Huynh, T.N., Jurado, A.R., Goo, Y.A., Sadilek, M., Iavarone, A.T., Sauer, J.D., Tong, L., and Woodward, J.J. (2014) The cyclic dinucleotide c-di-AMP is an allosteric regulator of metabolic enzyme function. Cell 158: 1389–1401.

Teh, W.K., Dramsi, S., Tolker-Nielsen, T., Yang, L., and Givskov, M. (2019) Increased Intracellular Cyclic di-AMP Levels Sensitize *Streptococcus gallolyticus* subsp. gallolyticus to Osmotic Stress and Reduce Biofilm Formation and Adherence on Intestinal Cells. J Bacteriol 201: e00597–00518.

Tolner, B., Ubbink-Kok, T., Poolman, B., and Konings, W.N. (1995) Characterization of the proton/glutamate symport protein of *Bacillus subtilis* and its functional expression in *Escherichia coli*. J Bacteriol 177: 2863–2869.

Tosi, T., Hoshiga, F., Millership, C., Singh, R., Eldrid, C., Patin, D., Mengin-Lecreulx, D., Thalassinos, K., Freemont, P., and Gründling, A. (2019) Inhibition of the *Staphylococcus aureus* c-di-AMP cyclase DacA by direct interaction with the phosphoglucosamine mutase GlmM. PLoS Pathog 15: e1007537.

Underwood, A.J., Zhang, Y., Metzger, D.W., and Bai, G. (2014) Detection of cyclic di-AMP using a competitive ELISA with a unique pneumococcal cyclic di-AMP binding protein. J Microbiol Methods 107: 58–62.

Vitko, N.P., Spahich, N.A., and Richardson, A.R. (2015) Glycolytic dependency of high-level nitric oxide resistance and virulence in *Staphylococcus aureus*. MBio 6: e00045–00015.

Whiteley, A.T., Garelis, N.E., Peterson, B.N., Choi, P.H., Tong, L., Woodward, J.J., and Portnoy, D.A. (2017) c-di-AMP modulates *Listeria monocytogenes* central metabolism to regulate growth, antibiotic resistance and osmoregulation. Mol Microbiol 104: 212–233.

Whiteley, A.T., Pollock, A.J., and Portnoy, D.A. (2015) The PAMP c-di-AMP Is Essential for *Listeria monocytogenes* Growth in Rich but Not Minimal Media due to a Toxic Increase in (p)ppGpp. Cell Host Microbe 17: 788–798.

Wicke, D., Schulz, L.M., Lentes, S., Scholz, P., Poehlein, A., Gibhardt, J., Daniel, R., Ischebeck, T., and Commichau, F.M. (2019) Identification of the first glyphosate transporter by genomic adaptation. Environ Microbiol 21: 1287–1305.

Witte, C.E., Whiteley, A.T., Burke, T.P., Sauer, J.D., Portnoy, D.A., and Woodward, J.J. (2013) Cyclic di-AMP is critical for *Listeria monocytogenes* growth, cell wall homeostasis, and establishment of infection. MBio 4: e00282–00213.

Woodward, J.J., Iavarone, A.T., and Portnoy, D.A. (2010) c-di-AMP secreted by intracellular *Listeria monocytogenes* activates a host type I interferon response. Science 328: 1703–1705.

Zarrella, T.M., Metzger, D.W., and Bai, G. (2018) Stress Suppressor Screening Leads to Detection of Regulation of Cyclic di-AMP Homeostasis by a Trk Family Effector Protein in *Streptococcus pneumoniae*. J Bacteriol 200: e00045–00018.

Zeden, M.S., Schuster, C.F., Bowman, L., Zhong, Q., Williams, H.D., and Gründling, A. (2018) Cyclic di-adenosine monophosphate (c-di-AMP) is required for osmotic regulation in *Staphylococcus aureus* but dispensable for viability in anaerobic conditions. J Biol Chem 293: 3180–3200.

Zhu, B., and Stülke, J. (2018) SubtiWiki in 2018: from genes and proteins to functional network annotation of the model organism *Bacillus subtilis*. Nucleic Acids Res 46: D743–D748.

Zhu, Y., Pham, T.H., Nhiep, T.H., Vu, N.M., Marcellin, E., Chakrabortti, A., Wang, Y., Waanders, J., Lo, R., Huston, W.M., Bansal, N., Nielsen, L.K., Liang, Z.X., and Turner, M.S. (2016) Cyclic-di-AMP synthesis by the diadenylate cyclase CdaA is modulated by the peptidoglycan biosynthesis enzyme GlmM in *Lactococcus lactis*. Mol Microbiol 99: 1015–1027.

Zhu, Y., Xiong, Y.Q., Sadykov, M.R., Fey, P.D., Lei, M.G., Lee, C.Y., Bayer, A.S., and Somerville, G.A. (2009) Tricarboxylic acid cycle-dependent attenuation of *Staphylococcus aureus* in vivo virulence by selective inhibition of amino acid transport. Infect Immun 77: 4256–4264.

