## Supplemental Material for "Identification of the main glutamine and glutamate transporters in *Staphylococcus aureus* and their impact on c-di-AMP production"

### Supplementary material

**Table S1. Composition of the glucose defined media (GDM) used in this study**

| Ingredients | GDM | GDM+Gln | GDM+Glu | GDM+$NH_3$ | GDM+Gln+$NH_3$ | GDM+Glu+$NH_3$ |
|---|---|---|---|---|---|---|
| **Salts (g/L)** | | | | | | |
| KCl | 15 | 15 | 15 | 15 | 15 | 15 |
| NaCl | 47.5 | 47.5 | 47.5 | 47.5 | 47.5 | 47.5 |
| $MgSO_4$ 7H2O | 6.5 | 6.5 | 6.5 | 6.5 | 6.5 | 6.5 |
| (NH4)$_2$SO4 | 0 | 0 | 0 | 20 | 20 | 20 |
| Tris | 60.5 | 60.5 | 60.5 | 60.5 | 60.5 | 60.5 |
| **Carbon source (g/L)** | | | | | | |
| Glucose | 25 | 25 | 25 | 25 | 25 | 25 |
| **Amino acids (mg/L)** | | | | | | |
| L-Arg | 50 | 50 | 50 | 50 | 50 | 50 |
| L-Pro | 10 | 10 | 10 | 10 | 10 | 10 |
| L-Gln | 0 | 100 | 0 | 0 | 100 | 0 |
| L-Glu | 0 | 0 | 100 | 0 | 0 | 100 |
| L-Val | 80 | 80 | 80 | 80 | 80 | 80 |
| L-Thr | 30 | 30 | 30 | 30 | 30 | 30 |
| L-Phe | 40 | 40 | 40 | 40 | 40 | 40 |
| L-Leu | 90 | 90 | 90 | 90 | 90 | 90 |
| L-Gly | 50 | 50 | 50 | 50 | 50 | 50 |
| L-Ser | 30 | 30 | 30 | 30 | 30 | 30 |
| L-Asp | 90 | 90 | 90 | 90 | 90 | 90 |
| L-Lys | 50 | 50 | 50 | 50 | 50 | 50 |
| L-Ala | 60 | 60 | 60 | 60 | 60 | 60 |
| L-Trp | 10 | 10 | 10 | 10 | 10 | 10 |
| L-Met | 10 | 10 | 10 | 10 | 10 | 10 |
| L-His | 20 | 20 | 20 | 20 | 20 | 20 |
| L-Ile | 30 | 30 | 30 | 30 | 30 | 30 |
| L-Tyr | 50 | 50 | 50 | 50 | 50 | 50 |
| L-Cystine | 20 | 20 | 20 | 20 | 20 | 20 |
| **Vitamins (mg/L)** | | | | | | |
| Biotin | 0.1 | 0.1 | 0.1 | 0.1 | 0.1 | 0.1 |
| Thiamine | 2 | 2 | 2 | 2 | 2 | 2 |
| Nicotinic acid | 2 | 2 | 2 | 2 | 2 | 2 |
| Calcium pantothenate | 2 | 2 | 2 | 2 | 2 | 2 |
| **Metals (mg/L)** | | | | | | |
| $CaCl_2$ $2H_2O$ | 22 | 22 | 22 | 22 | 22 | 22 |
| $KH_2PO_4$ | 140 | 140 | 140 | 140 | 140 | 140 |
| $FeSO_4$ $7H_2O$ | 6 | 6 | 6 | 6 | 6 | 6 |
| $MnSO_4$ $H_2O$ | 7.58 | 7.58 | 7.58 | 7.58 | 7.58 | 7.58 |
| Citric acid | 6 | 6 | 6 | 6 | 6 | 6 |

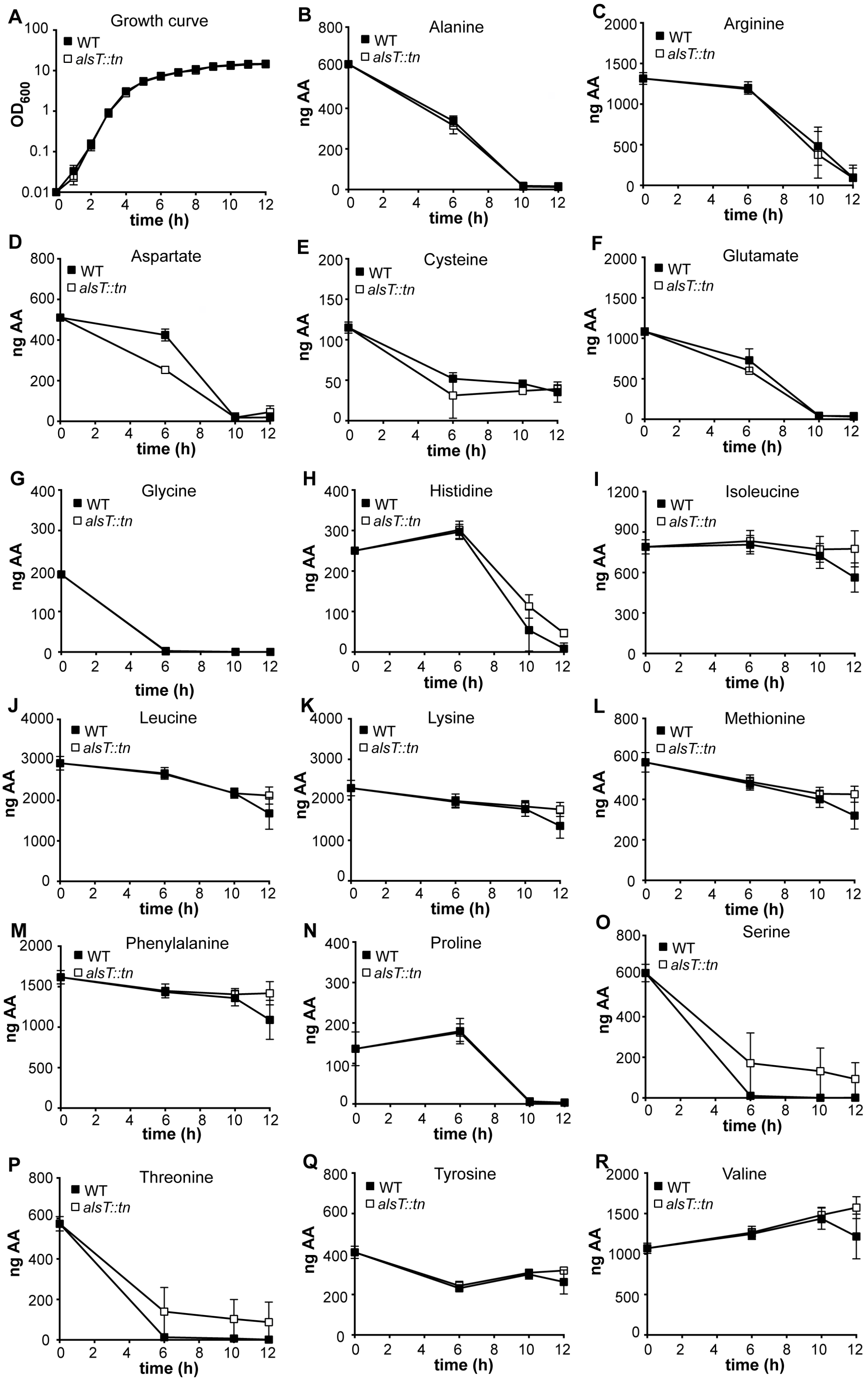
A Growth curve
B Alanine
C Arginine
D Aspartate
E Cysteine
F Glutamate
G Glycine
H Histidine
I Isoleucine
J Leucine
K Lysine
L Methionine
M Phenylalanine
N Proline
O Serine
P Threonine
Q Tyrosine
R Valine
WT
alsT::tn
OD600
ng AA
time (h)

**Fig. S1: Growth and amino acid consumption analysis to determine the function of AlsT.** (A) Bacterial growth curves. *S. aureus* strains LAC* (WT) and LAC**alsT::tn* (*alsT::tn*) were grown in TSB medium and $OD_{600}$ readings determined at hourly intervals and the average and SDs from three biological replicates plotted. (B-R) Quantification of amino acid levels in culture supernatants. Spent medium samples from the cultures shown in panel A were prepared at the 0, 6, 10 and 12 h time points and the amino acid level determined as previously described using an amino acid analyzer (Halsey *et al.*, 2017). Amino acids analyzed are indicated above each panel. The average values and SDs from three biological replicates were plotted.

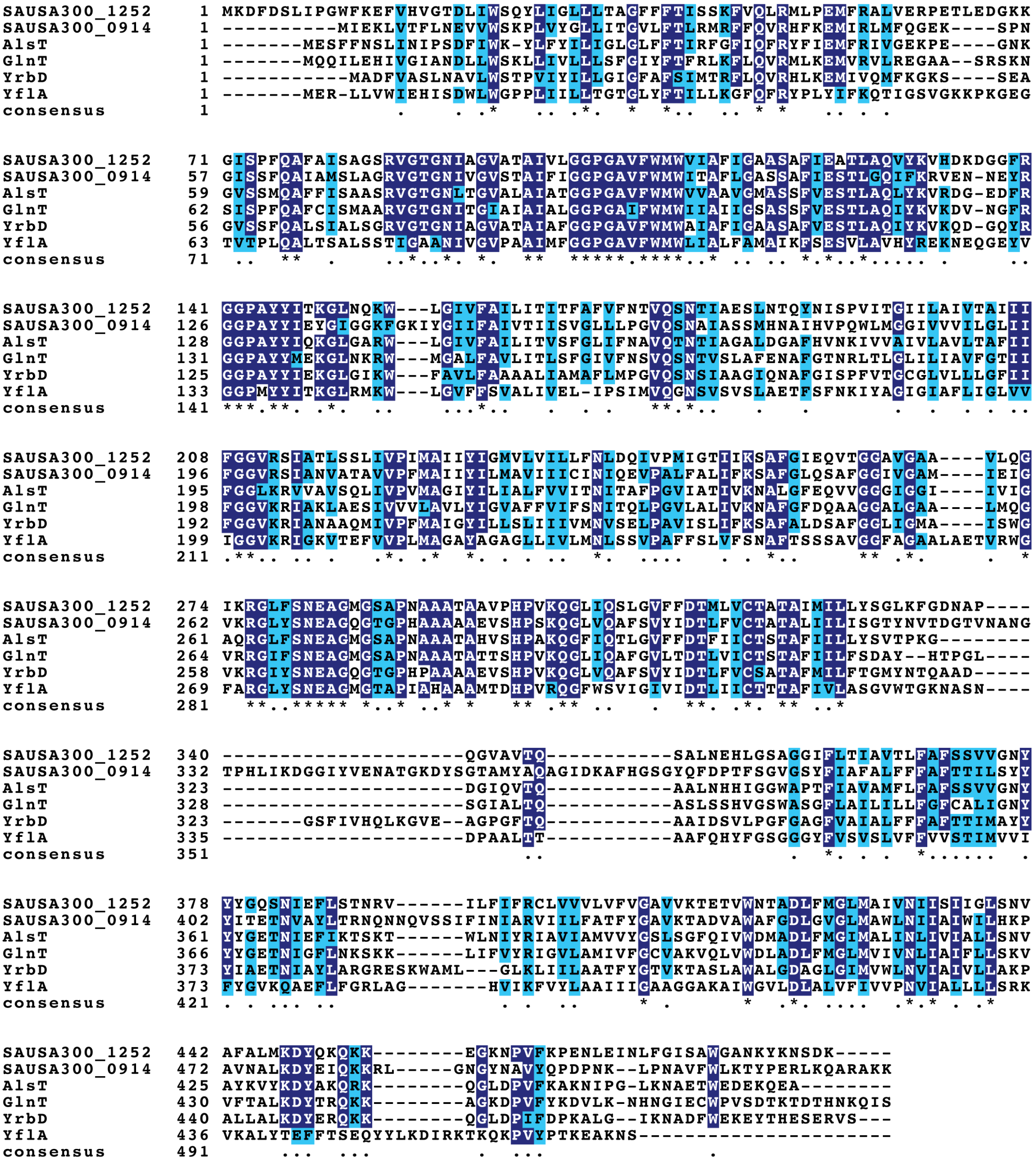

**Fig. S2. CLUSTAL Omega alignment of *B. subtilis* and *S. aureus* AlsT/GlnT homologs.** The *S. aureus* AlsT (SAUSA300_1252) protein was used as query sequence in a BLASTP search to identify homologs encoded in *S. aureus* FPR3757 and *B. subtilis* 168. Besides, AlsT (SAUSA300_1252), one additional homolog, SAUSA300_0914, was identified in *S. aureus* and four close homologs $AlsT_{BS}$, $GlnT_{BS}$, $YrbD_{BS}$ and $YflA_{BS}$ were found in *B. subtilis.* A Clustal-Omega alignment was performed with the six proteins and identical residues were shaded in dark blue and similar residues in light blue.

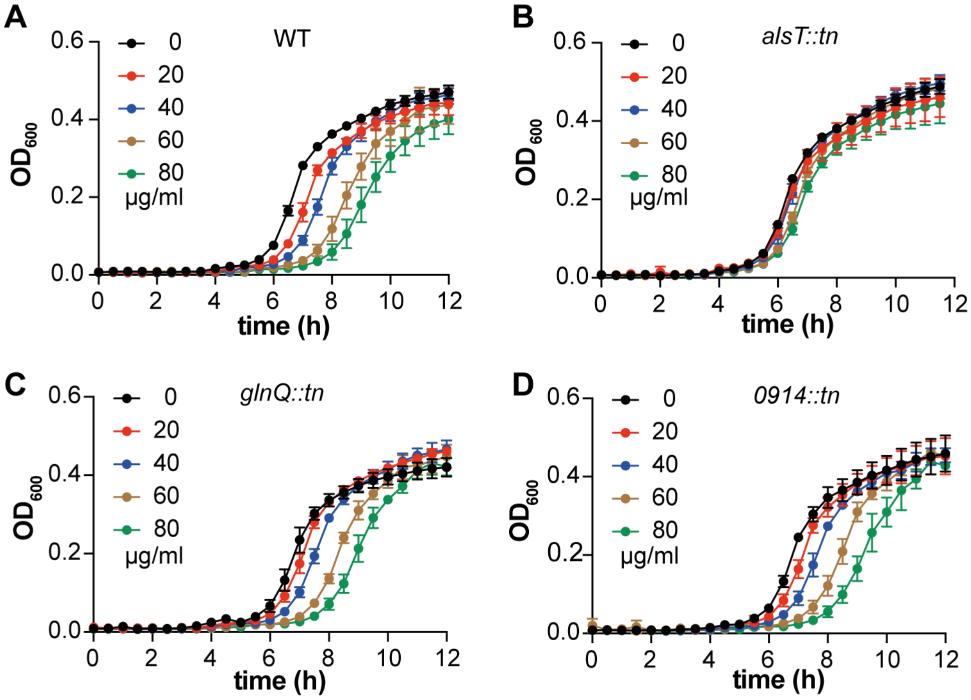


**Fig. S3. LAC*_alsT::tn_ shows increased resistance to the toxic glutamine analogue γ-L-glutamyl hydrazide.** (A-D). Bacterial growth curves. *S. aureus* strains (A) LAC* (WT), (B) LAC*_alsT::tn_ (*alsT::tn*), (C) LAC*_glnQ::tn_ (*glnQ::tn*) and (D) LAC*_0914::tn_ (*0914::tn*) were grown for 12 h in GDM+$NH_3$ in the presence of 0 (black), 20 (red), 40 (blue), 60 (brown) and 80 (green) µg/ml of γ-L-glutamyl hydrazide. Average $OD_{600}$ values and SDs of three independent biological replicates were plotted.

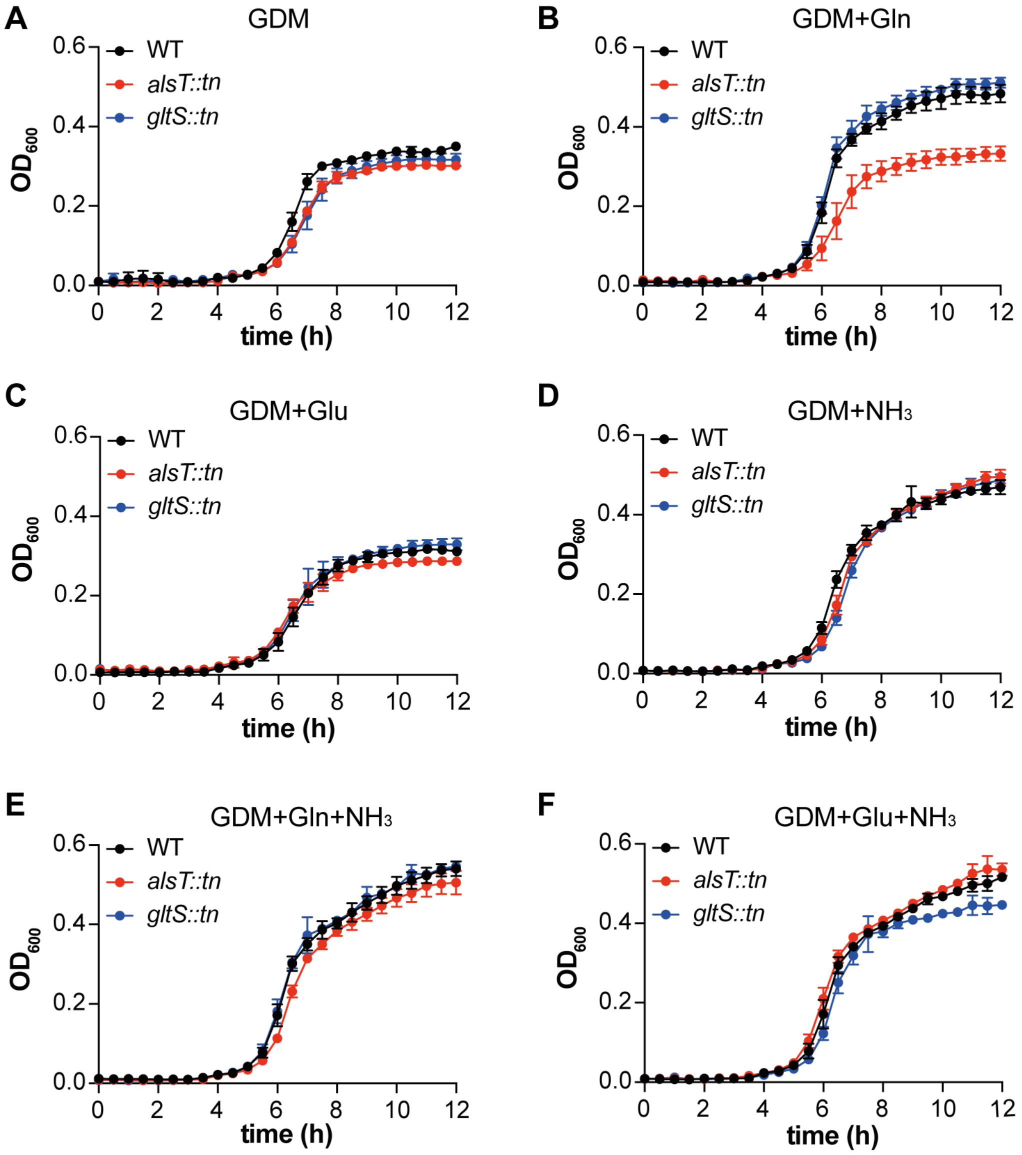


**Fig. S4. Lack of glutamine uptake results in growth deficiency of *S. aureus* grown in GDM+Gln.** (A-F) Bacterial growth curves. *S. aureus* strains LAC* (WT), LAC**gltS::tn* (*gltS::tn*) and LAC**alsT::tn* (*alsT::tn*) were grown in either (A) GDM, (B) GDM+Gln, (C) GDM+Glu, (D) GDM+$NH_3$, (E) GDM+Gln+$NH_3$, or (F) GDM+Glu+$NH_3$ for 12 h and $OD_{600}$ readings determined. Average $OD_{600}$ readings and SDs from three biological replicates were plotted.

**REFERENCES:**

Halsey, C.R., Lei, S., Wax, J.K., Lehman, M.K., Nuxoll, A.S., Steinke, L., Sadykov, M., Powers, R., and Fey, P.D. (2017) Amino Acid Catabolism in *Staphylococcus aureus* and the Function of Carbon Catabolite Repression. *MBio* **8**: e01434-01416.
